# Deep Insertion, Deletion, and Missense Mutation Libraries for Exploring Protein Variation in Evolution, Disease, and Biology

**DOI:** 10.1101/2022.07.26.501589

**Authors:** Christian B. Macdonald, David Nedrud, Patrick Rockefeller Grimes, Donovan Trinidad, James S. Fraser, Willow Coyote-Maestas

**Affiliations:** Department of Bioengineering and Therapeutic Sciences, University of California, San Francisco, United States; Bio-Techne, Minneapolis, Minnesota, United Stated; Department of Medicine, Division of Infectious Disease, University of California, San Francisco, United States; Quantitative Biosciences Institute, University of California, San Francisco, United States

## Abstract

Insertions and deletions (indels) are a major source of genetic variation in evolution and the cause of nearly 30% of Mendelian disease. Despite their importance, indels are left out of nearly every systematic mutational scan to date due to technical challenges associated with making indel-containing libraries, limiting our understanding of indels in disease, biology, and evolution. Here we present a library generation method, DIMPLE, that generates deletions, insertions, and missense at similar frequencies within any gene. To benchmark DIMPLE, we generated libraries within four genes (Kir2.1, VatD, TRPV1, and OPRM1) of varying length and evolutionary origin. DIMPLE produces libraries that are near complete, low cost, and low bias. We measured how missense mutations and indels of varying length impact the potassium channel Kir2.1 surface expression. Across all Kir2.1’s secondary structure, deletions are more disruptive than insertions, beta sheets are extremely sensitive to large deletions, and flexible loops allow insertions far more frequently than deletions. DIMPLE’s low bias, ease of use, and low cost will enable high throughput probing of the importance of indels in disease and evolution.

## Introduction

Mutations are one of the fundamental tools biologists use to understand the nature of genes. To understand how proteins work, biochemists mutate amino acids to learn which are important. Evolutionary biologists reconstruct the history of changes in a gene to understand how its function changes over time. Synthetic biologists create improved enzymes by introducing mutations and screening for catalytic improvement. Clinical geneticists infer pathogenicity using machine learning that integrates systematic mutational scanning data, conservation patterns, and variant frequencies within patient populations. Each paradigm has produced fundamental insights into how nature produces life and what goes wrong in disease, but each often overlooks mutations beyond simple substitutions. Recent work has underscored how essential other types of mutations are to evolutionary novelty and adaptation, as well as their utility for understanding diseases and protein engineering (Seuma, Lehner, and Bolognesi 2022; Savino, Desmet, and Franceus 2022; Q. Ma et al. 2022; Park and Hahn 2021; Z. Zhang et al. 2018; Ogden et al. 2019). To evaluate how mutations change proteins, in addition to missense mutations we must consider frameshifts, recombination, splice variations, and insertions and deletions. Non-missense mutations present challenges for sequence alignment and evolutionary models and the lack of a biophysical model for how they impact proteins limits their use by protein engineers and understanding by biologists.

Massively-parallel mutational scanning, where systematic sets of mutations are created and then profiled by selection or screening, is commonly used to understand the nature of changes in a protein sequence. Mutational scanning has a long history in experimental biology, starting from pre-molecular techniques such as random cloning for gene mapping (Kohara, Akiyama, and Isono 1987). Improved enzymes and sequencing allowed site-directed mutagenesis and iterative small-scale cysteine and alanine scans (Morrison and Weiss 2001; Zhu and Casey 2007). These require iterative mutagenesis and verification for each variant, making them labor-intensive. Error-prone PCR offers simpler access to libraries of mutant sequences, but it is not programmable and not systematic (which may be of benefit for directed evolution efforts) (Drummond et al. 2005). Systematic variant libraries were enabled using parallel inverse PCR coupled to degenerate codons which enabled reliable variant libraries, but library composition was thus limited to a handful of degenerate codon schemes (Pines et al. 2015; Hughes et al. 2003). Inverse PCR for creating genotypic diversity coupled with sequencing-based assays for phenotypes form the basis for Deep Mutational Scanning (DMS) (Fowler and Fields 2014). DMS studies are enabling fundamental insights in protein biochemistry, evolution, and the molecular basis of disease. These efforts have culminated in large-scale international efforts such as the Atlas of Variant Effects Alliance, with the goal of characterizing all variants circulating within human populations. However, while insertions and deletions (indels) make-up nearly ⅓ of disease-causing variants, to date only one pioneering DMS study on disease causing genes has included indels (Seuma, Lehner, and Bolognesi 2022).

DMS studies do not include indels primarily because most DMS libraries are constructed using inverse PCR. While inverse PCR works well for missense variant libraries to make deletion or insertions variants, individual primers would be needed for every variant. For this reason transposons are most commonly used for indel library generation (Emond et al. 2020; Edwards et al. 2008; Liu et al. 2016). However due to bias intrinsic to transposons these libraries are incomplete, imbalanced, and do not work well for some targets (Green et al. 2012; Coyote-Maestas et al. 2020). An alternative to inverse-PCR and transposon based approaches is to leverage microarray-based oligo synthesis (OLS) for making systematic mutational libraries (Kitzman et al. 2015; Kowalsky et al. 2015; Melnikov et al. 2014). The basic principle of these approaches is to synthesize the variants of interest within all positions of a subregion of a gene and stitch in mutated subregions by recombination or restriction-ligation cloning. Because each variant is individually synthesized rather than randomly generated, OLS-based libraries are typically more complete, can include any variant type, and are simpler to clone than PCR-based approaches. Indeed, the only two mutational scans to date that included indel variants, Seuma et al and Ogden et al. 2019, were made using an OLS-based approach.

Here we present a combined design and experimental pipeline, Deep Indel Missense Programmable Library Engineering (DIMPLE), based on OLS-based synthesis and golden gate cloning. DIMPLE consists of a solution for library design, synthesis, and quality control. Our libraries are an improvement in complexity, completeness, bias and affordability compared to previous methods. To demonstrate the utility of DIMPLE, we apply it to study how indels impact surface expression of the model potassium channel Kir2.1. This dataset is the first systematic indel scan within a large multi-domain protein, which allows us to empirically explore how insertions and deletions impact protein structure. We compare our data to variants present in the clinic and homologous proteins to explore indels in inward rectifier disease and evolution.

### A method to generate libraries containing point, insertion, and deletion mutations in parallel

We designed the DIMPLE pipeline to produce libraries with deletions and insertions along with point mutations at all positions of a gene. We built DIMPLE using a previous library generation pipeline, SPINE, as a scaffold which we developed for domain insertion scanning and later extended for missense mutational scanning (Coyote-Maestas et al. 2020; Nedrud, Coyote-Maestas, and Schmidt 2021). DIMPLE encodes mutational diversity in microarray-based oligo pools which are assembled together with final libraries with PCR amplified fragments containing the remainder of the gene and its vector (Fig 1A). By using oligo pools, mutational diversity is precisely controlled, and assembly removes bias that occurs in inverse-PCR and transposon-based libraries. The DIMPLE software automates the process by generating primers for amplifying sublibraries, primers for amplifying the backbone and adding complementary golden-gate compatible cutsites, and mutated oligo pools (https://github.com/coywil26/DIMPLE, Supp Fig 1A). To assist the community in making DIMPLE libraries, we wrote a detailed open-source protocol deposited on protocol.io: (https://dx.doi.org/10.17504/protocols.io.rm7vzy7k8lx1/v1).

**Figure 1.**
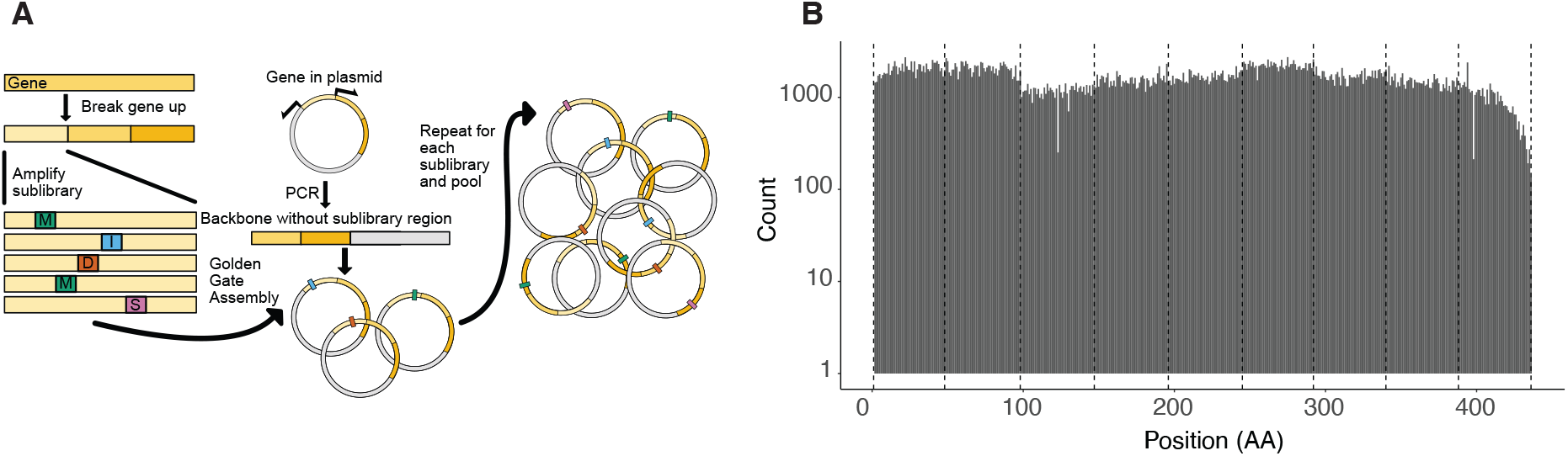
Generation of programmed mutational, deletional, and insertional libraries with DIMPLE in the model potassium channel Kir2.1. **a**. Schematic depiction of the library generation process with DIMPLE. **b**. Barplot of mutation type per position against counts. All variants are stacked. The boundaries of each mutagenic sublibrary are indicated with dashed lines. Overall, each mutagenized sublibrary region is within 2-fold of eachother implying well-balanced libraries (supplemental figure 3B).

For DIMPLE to be useful for the scientific community at large, it should work on as broad a range of targets as possible. To test whether the computational portion of the pipeline can generate variants in genes spanning a range of length and composition, we tested it against 24 genes, ranging in length from 42 to 2561 amino acids and 43% to 59% GC content, yielding 279 fragments and 395330 total variants (Supp. Table 1). In all cases in-silico assembly succeeded in yielding in-frame assemblies with all expected variants present.

When considering the lengths of insertions and deletions to include in our libraries, we wanted to balance the number of variants and library size with the potential for insight and specificity. The length distribution of deletions in human genomes follows a power law, which suggests that larger deletions will be exponentially rare (J. Zhang et al. 2010). The one prior indel-scanning experiment revealed that increasingly-long indels are highly system-specific, with idiosyncratic effects of large deletions being driven by exposure of a particular nucleating core and unlikely to generalize (Seuma, Lehner, and Bolognesi 2022). We chose 1-3 amino acid long indels for our default libraries, as this allowed us to capture most relevant variation and potentially observe any non-linear effects while maximizing sequencing capacity. DIMPLE is ultimately constrained by oligo synthesis, however, and the Agilent 230 bp platform we use would allow systematic screens up to 27 bp deletions and 120 bp insertions.

As a demonstration of DIMPLE’s utility, we generated a library with the potassium channel Kir2.1 which contains at every amino acid a mutation to every other amino acid and when possible, a synonymous mutation, 1-3 codon deletions, and 1-3 codon insertions (G, GS, GSG), thus 26 variants per residue. We integrated these libraries into stable cell lines using a commonly used high efficiency landing pad cell line method optimized for library generation (Matreyek et al. 2020).

DIMPLE is an easy-to-use and customizable computational and experimental pipeline with thorough documentation for generating effective mutational libraries with diverse variant types.

### DIMPLE libraries have even coverage across positions, variant types, and gene targets

Mutational scanning experiments are critically dependent on library quality. In DMS screens we measure a change in frequency over time, meaning any over or underrepresented variants in a starting library will decrease assay sensitivity and introduce noise. An ideal library generation method should reliably produce variant pools with even representation a) across variants at each position, b) between positions across the target, c) have nearly all variants present, and d) be target gene agnostic.

With DIMPLE, we attempted to meet these goals for substitutions, insertions, and deletions. Indels introduce an additional difficulty, as they alter the overall length of synthesized oligos. In indel libraries, each mutagenic region consists of a range of sizes. We worried this would introduce bias during sublibrary PCR amplification, yielding systematic bias between variant types. To avoid this, we include buffer sequences outside the Golden Gate cut sites for deletion and substitution variants which are adjusted for each variant type, keeping all oligos within a sub-library the same length during amplification but allowing a range of sizes after assembly (Supp Fig 2). To test if different variants are present at similar frequencies, we compared the distributions of each variant across the entire gene. We find most variants are present at similar frequencies, however in Kir2.1 it appears that most indels are present at slight yet significantly reduced frequencies. That said, there is less than a two-fold difference between all variant types (Figure 2A).

**Figure 2.**
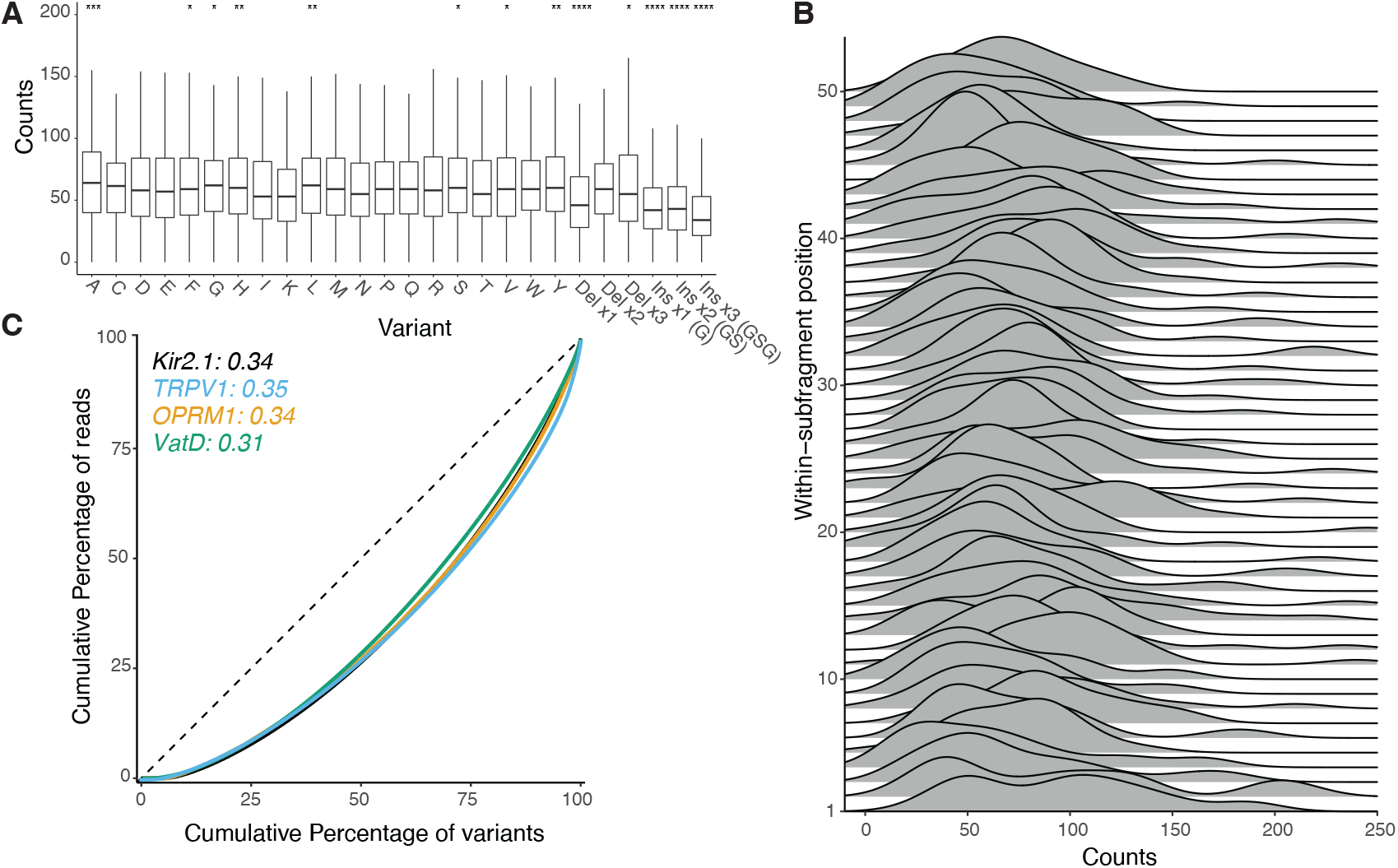
Quantifying the bias of library assembly with DIMPLE. **A**. Boxplots of variants at each position across all of Kir2.1. The vertical length of the box is the interquartile range (IQR), upper bound is the 75th percentile with the lower bound is the 25th percentile. Significance is tested using two-sided t-tests controlled for multiple comparisons comparing incorporation means between variants across all positions. Significance levels: ***P < 0.001; **P < 0.01; *P < 0.05, all others not significant. **B**. Stacked density plots, or ridge plot ordered bottom-to-top from first to last positions of the second sublibrary of Kir2.1. **C**. Lorentz curves and Gini coefficients test the inequality within the distribution of observed variants. A completely even distribution would be a diagonal with a Gini score of 0. The distribution of designed variants for mutagenic libraries of Kir2.1, TRPV1, VatD, and OPRM1 are shown with corresponding Gini scores noted.

Positional bias is a frequent problem and challenge for DMS libraries. For microarray-based library generation methods which require manual sublibrary pooling step, the largest source of positional variability comes from this mixing. This is apparent by eye, with the sublibrary three having the lowest and four having the highest frequency Figure 1B). We find across Kir2.1, that median mutational frequencies across sublibraries are all within 2-fold, implying we have evenly represented libraries (Figure 1B, Supp. Fig 3).

In previous oligo-pool derived libraries, we observed low mutation rates at the beginnings and ends of sublibraries compared to the middle (Coyote-Maestas et al. 2020). We wondered if this might reflect a positional effect on digestion or ligation efficiency. To address this potential source of bias, DIMPLE includes 4 non mutated residues from the wildtype sequence immediately after and before the first and second cutsites, respectively. We tested the impact of this modification by comparing the within-sublibrary distributions of variants in each sublibrary (Figure 2B, Supp Fig 3). We find no systematic positional biases within sublibraries, implying low bias in each oligo pool.

To test the robustness of our technique across different targets from a variety of organisms and classes, we generated additional libraries of a bacterial antibiotic resistance element (VatD from *Enterococcus faecium*), the rat temperature-sensing ion channel TRPV1, and human μ-opioid receptor OPRM1. As with Kir2.1, these libraries contain nearly every variant (VatD-97.5%, TrpV1-97%, and 93.2% out of 5408, 21754, and 10412 possible variants, respectively), with representation at similar frequencies positionally across all sublibraries within two-fold of the mean, within two-fold by variant types across positions, and similar variant incorporation at positions within sublibraries (Fig 2C, Supp Fig 4-5). We are confident therefore that DIMPLE succeeds at generating missense, insertion, and deletional variants across a range of targets.

In summary, DIMPLE generates libraries that are affordable (<0.30$/variant, Supp. Table 2), near complete, with little bias across positions and variant types, and robust to different targets.

### DIMPLE libraries allow access to unexplored sequence space, revealing how indels impact Kir2.1 surface expression

Our initial target, Kir2.1, is a potassium channel with a variety of physiological roles, primarily setting the resting membrane potential of a cell (Hager et al. 2021). Many mutations, including deletions, impact Kir2.1 surface expression and cause severe cardiac and developmental disorders (Hager et al. 2021; Donghui Ma et al. 2007). To understand how indels affect Kir2.1 physiology, we performed an assay to specifically identify mutational impacts on surface trafficking. We generated stable cell lines with our Kir2.1 DIMPLE libraries in HEK293T cells, sorted the Kir2.1 DIMPLE libraries based on specific Kir2.1 surface expression with a fluorescent antibody into subpopulations, then sequenced these populations to determine the genotype of variants within each population. By calculating enrichment of variants across surface expression populations relative to WT Kir2.1, we determine how 10964 (out of a total possible 11302, or 97%) variants impact surface expression (Fig 3B, Supp Fig 6). For these fitness scores, we find high reproducibility between three replicates and our previous study with missense mutations that used the same surface expression assay (Supp Fig 7). Across this dataset, we see a clear hierarchy of impacts across mutation types, with (on average) missense mutations being more harmful than synonymous mutations, insertions much more pronounced, and most deletions deleterious for trafficking (Fig 3C). The distribution of fitness effects here appears bimodal, with a population of WT-like variants, with a long tail of rare improved-trafficking variants, and a second population of poorly trafficked variants. Synonymous mutations preserve the protein sequence, and so their influences would be limited to second-order effects translation and/or transcription. As expected, these variants are unimodal, centered around neutrality. Substitutions are extremely context-dependent, with the impact of each depending on the physicochemical context in the structure. Consistent with other indel mutagenesis studies, we observed that deletions were in general more deleterious than insertions (Ogden et al. 2019; Gonzalez, Roberts, and Ostermeier 2019; Seuma, Lehner, and Bolognesi 2022; Arpino et al. 2014). This effect becomes stronger with increasing length for both insertions and deletions as well (Supp Fig 8).

**Figure 3.**
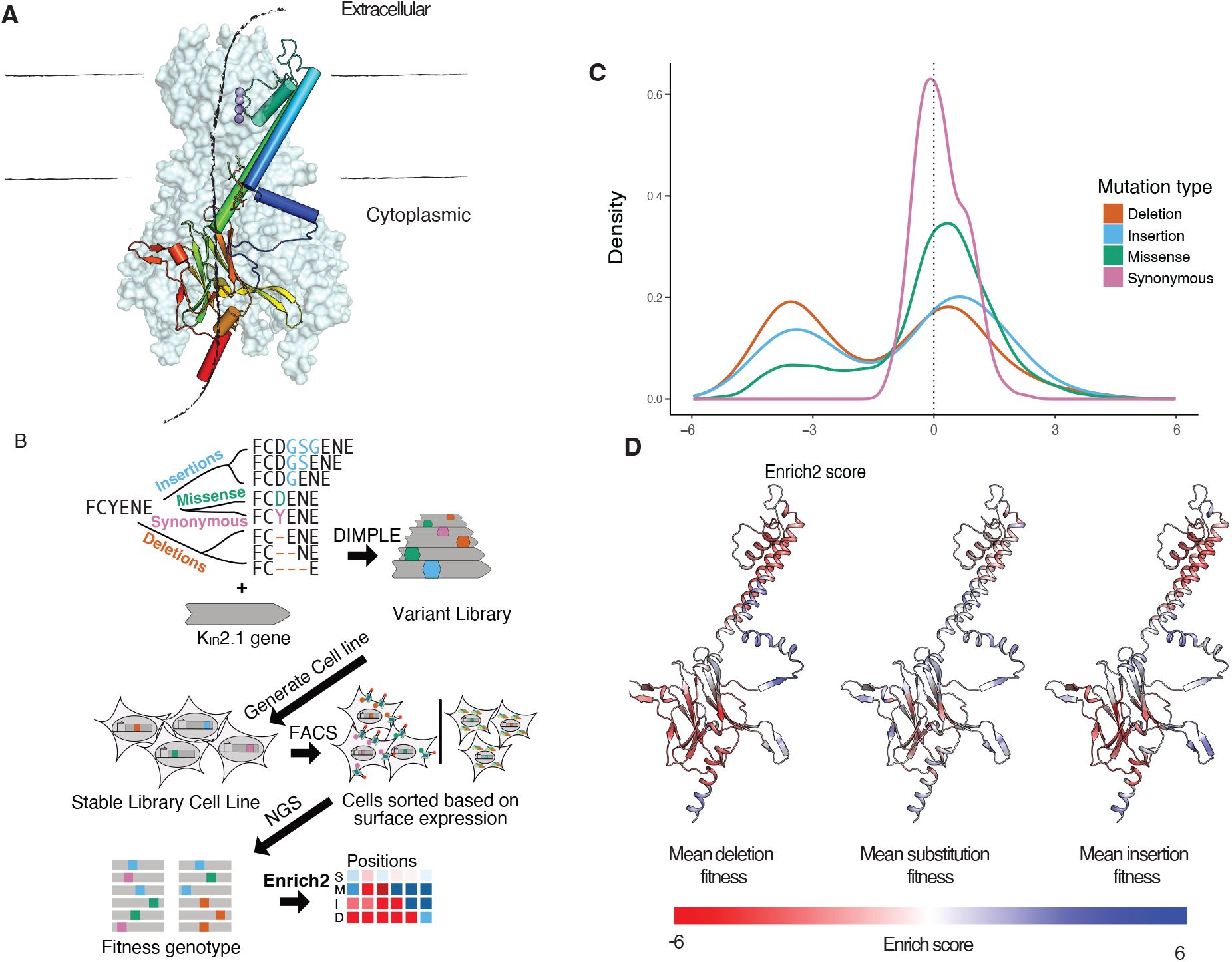
Variable-length indel scanning of Kir2.1 membrane trafficking. **a**. Cartoon schematic of Kir architecture: the monomeric structure and overall tetrameric assembly are shown with the crystal structure of Kir2.2 (3SPI). Boundaries of the lipid membrane are indicated with lines, the crystallographic potassium are shown in purple, and locations of the pore highlighted with a cartoon arrow crossing through the channel (Lomize et al. 2012). **b**. Cartoon workflow for studying how different variant types impact Kir2.1 surface expression. Briefly, we use DIMPLE to generate a library including insertion, missense, synonymous, and deletion variants at all positions of Kir2.1, we generate stable HEK293 cell lines, sort these cells based on surface-expression using FACS, sequence these subpopulations using Illumina Novaseq, and calculate surface expression fitness scores using Enrich2. **c**. The distribution of fitness effects on surface expression of Kir2.1 is displayed as a kernel density estimate. Negative scores indicate decreased trafficking relative to WT Kir2.1. Deletions are the most disruptive perturbation, followed by insertion, missense, and synonymous mutations, respectively. **d-f**. Mapping the average fitness effects of deletions, substitutions, and insertions across homologous positions in Kir2.2 shows global similarities but local differences between perturbation types. These are plotted from red-white-blue based on surface fitness scores. In general, the structured regions of Kir2.1 are more sensitive to all mutation types.

Examining the pattern of mutational effects on Kir2.1, we found many regions where effects on trafficking were similar across all variant types (Figs 3D and 4, Supp Fig 9). In some cases, the reasons are obvious, such as the FLAG tag (positions 116-123) where mutations disrupt antibody labeling. Across all mutation types, the unstructured N and C termini (positions 1-55 and 378-442) are more mutable than structured regions. Similarly, several flexible loops, such as the βE-βG and βH-βI loops, tolerate any mutations. In the helical (e.g., H109-L112 and V130-Q147) that determines potassium channel folding, and folding critical regions of the cytosolic C-terminal domain are completely immutable (e.g.,F203-V221, T276-D289, and S322-Y334) (Gajewski et al. 2011; Fallen et al. 2009). Overall, as in our previous DMS of Kir2.1 secondary structural elements are less mutable than unstructured regions (Coyote-Maestas et al. 2022) (Figs 3D and 4).

**Figure 4.**
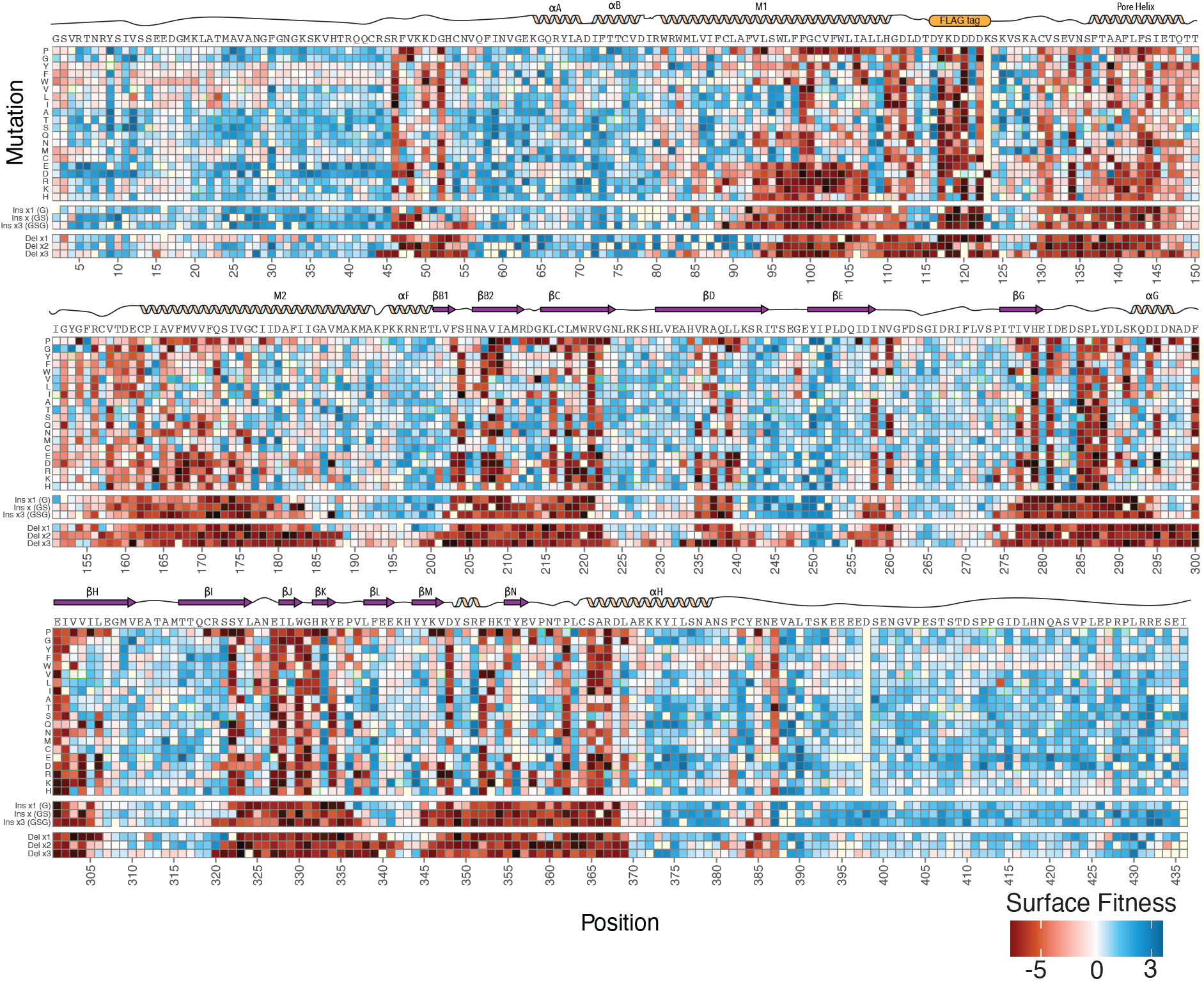
Mutational scanning shows the structural logic of trafficking in Kir2.1. Heatmap of surface expression fitness scores calculated from Enrich2 gradient colored from red (less than WT fitness) to white (WT fitness) to red (greater than WT fitness). Cartoon of secondary structure and labels of structural elements denoted above the heatmap. Only positions for which there were reads in all three replicates are shown here; others were removed in enrichment calculations. Synonymous mutation boxes are outlined with green. Mutations without data are highlighted in light yellow

Deletions are commonly used by biochemists in an *ad hoc* fashion to find and test important motifs. For example, Lily Jan’s group identified two motifs within Kir2.1 that were necessary for cell surface expression, the FCYENE and SY motifs (Dzwokai Ma et al. 2001). The FCYENE is a classic example of a diacidic ER-export motif. While the SY motif was later determined to be a golgi export motif that is a binding interface for the trafficking pathway component, AP1 adaptor γ protein (Donghui Ma et al. 2011). Deletions within the SY motif are extremely deleterious in our assay, while the FCYENE motif deletions are moderately disruptive. The FCYENE is in the distal C terminus in non-folding critical regions meaning mutations here likely solely impact ER-export. In contrast, the SY motif interacts directly with the hydrophobic core so SY variants will additionally suffer dramatic folding deficits (Donghui Ma et al. 2011; Coyote-Maestas et al. 2022; Li et al. 2016). With DIMPLE, we can confirm existing phenotypes within known trafficking motifs and in less understood proteins could discover new trafficking motifs and their boundaries.

### Insertions and deletions have distinct impacts dependent on Kir2.1 secondary structure

The impact of missense mutations within secondary elements, appear dependent on the physicochemistry of the mutation. In contrast, indels are broadly disruptive within secondary structural elements enabling a form of secondary structure footprinting. Despite similarities broadly across secondary structures, insertions, deletions, and varying lengths distinctly impact Kir2.1 surface expression. For example, within the αA and αB slide helices, deletions are generally beneficial with larger ones being most beneficial. The slide helix undergoes a disorder-to-order transition upon ligand so perhaps removing this semi-disordered region provides a stabilizing impact on folding perhaps reflecting a tradeoff between folding stability and function (Figure 5A-B). In contrast, within M1 deletions 1-2 AA deletions are beneficial for surface expression whereas 3 AA deletions and all insertions are neutral or deleterious (Supp Fig 10). It is hard to intuit exactly what is occurring but perhaps because M1 pivots in channel opening, it could have additional slack that deletions are removing. Due to M1’s role in function, we anticipate these variants are non-functional. To explore how insertions and deletions impact beta sheets we compared how different length insertions and deletions impact βD, βH, and βI (Figure 5C-D). βD and βH are for the most part completely intolerant to indels. βI in contrast is surprisingly tolerant to deletions, with the entirety of the beta sheet allowing 1 AA deletions and 2-3 AA deletions allowed in most of the beginning. In βI, G and GS insertions appear to be somewhat tolerated with GSG insertions quite deleterious throughout. βI does not appear to be entirely necessary for folding whereas βD and βH are. While overall indels within secondary structure elements are disruptive, within alpha helices and beta sheets there are surprising differences in sensitivity between indels with varied lengths. Within Kir2.1 the beta sheets overall appear to be far more sensitive to insertions and deletions than alpha helices.

**Figure 5.**
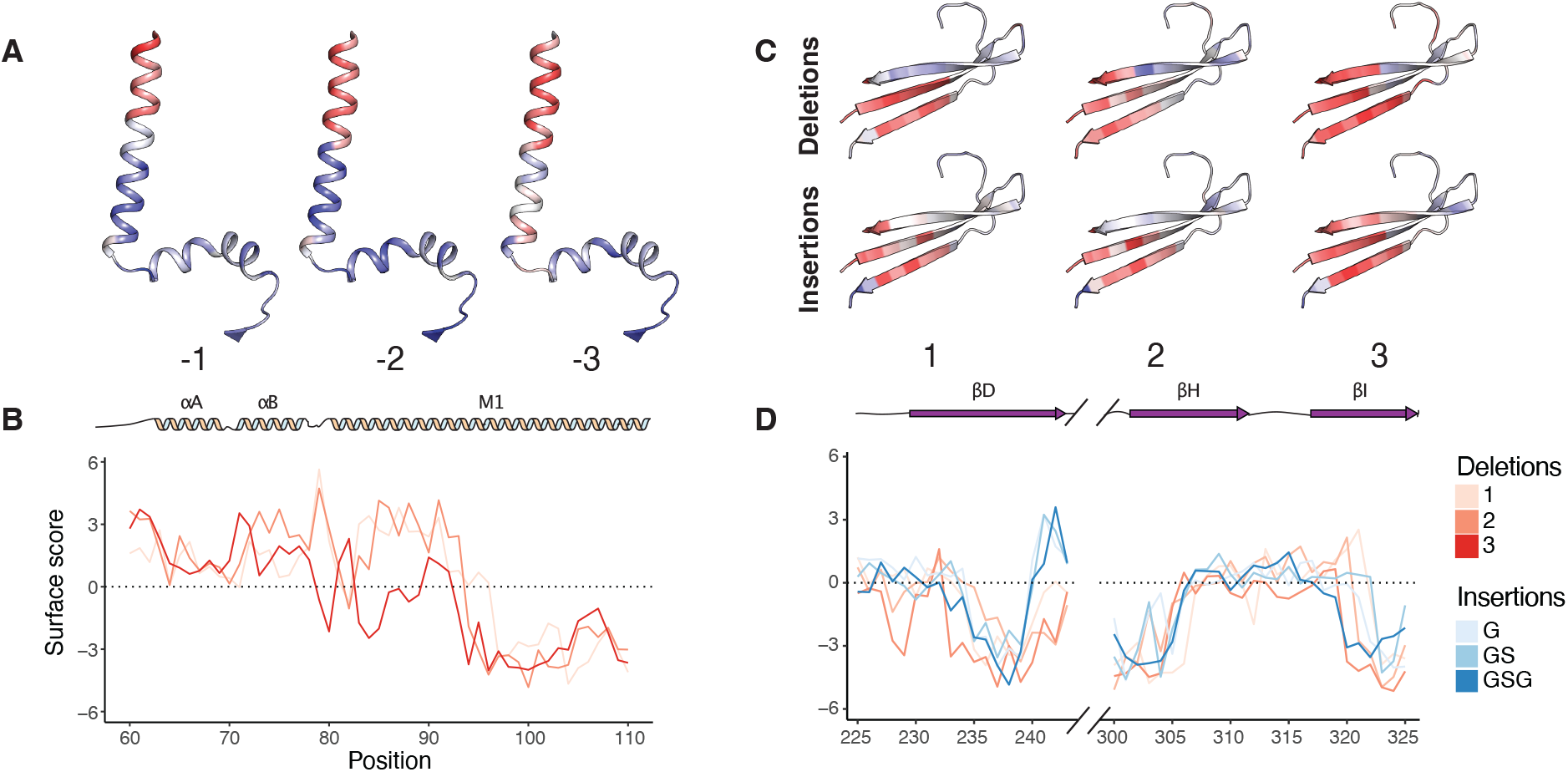
The length of an insertion and deletion impacts Kir2.1 surface expression. **a**. Impact of varying the length of deletion on surface expression mapped onto the M1 transmembrane alpha helix and slide helix colored from low-to-high surface expression, red-to-white-to-blue, respectively. **b**. Surface scores for the slide-helix position with varying lengths of deletions colored with increasing hue for increasing (or decreasing) length. 1-2 amino acid deletions are tolerated while 3 amino acids result in substantially less surface expression. **c**. Impact of varying the length of deletion (top) and insertion (bottom) on surface expression mapped onto three of the immunoglobulin beta sheets colored from low-to-high surface expression, red-to-white-to-blue, respectively. **d**. Surface scores for the beta sheet position with varying lengths of deletions colored with increasing hue for increasing (or decreasing) length. Because one of the beta sheets is not concurrent with the others only the beta sheets are focused on within the line plot.

### Insertions and deletion in disease and evolution

Insertions and deletions play a major role in disease. On average ⅔ of these will also cause a frameshift, unsurprisingly causing major disruptions. There are several important examples of in-frame deletions being associated with disorders, including Δ508 in CFTR (Lukacs and Verkman 2012). There is evidence for the pathogenicity of two deletions in Kir2.1 (ΔA91-L94 and ΔS314-Y315) and two additional deletions are of unknown significance (ΔA306 and ΔF99) (Landrum et al. 2018). While ΔA91-L94 is not contained within our library because it is four AA long, both ΔA91-ΔA93 and ΔA92-ΔA94 have extremely low surface fitness scores (Fig 6B). Putative pathogenic mutation ΔS314-Y315 and variant of unknown significance (VUS) ΔF99 are both within folding critical regions and have extremely low surface fitness scores and so are likely pathogenic. The VUS ΔA306 is unambiguously neutral in our data despite being in the gLoop, which is critical for potassium conductance. Our fitness data is based on an assay for surface expression if we added an additional screen for function ΔA306 would likely be functionally disruptive and potentially pathogenic. Indel scanning helps us explore the molecular mechanisms and potential pathogenicity of indels in human disease.

**Figure 6.**
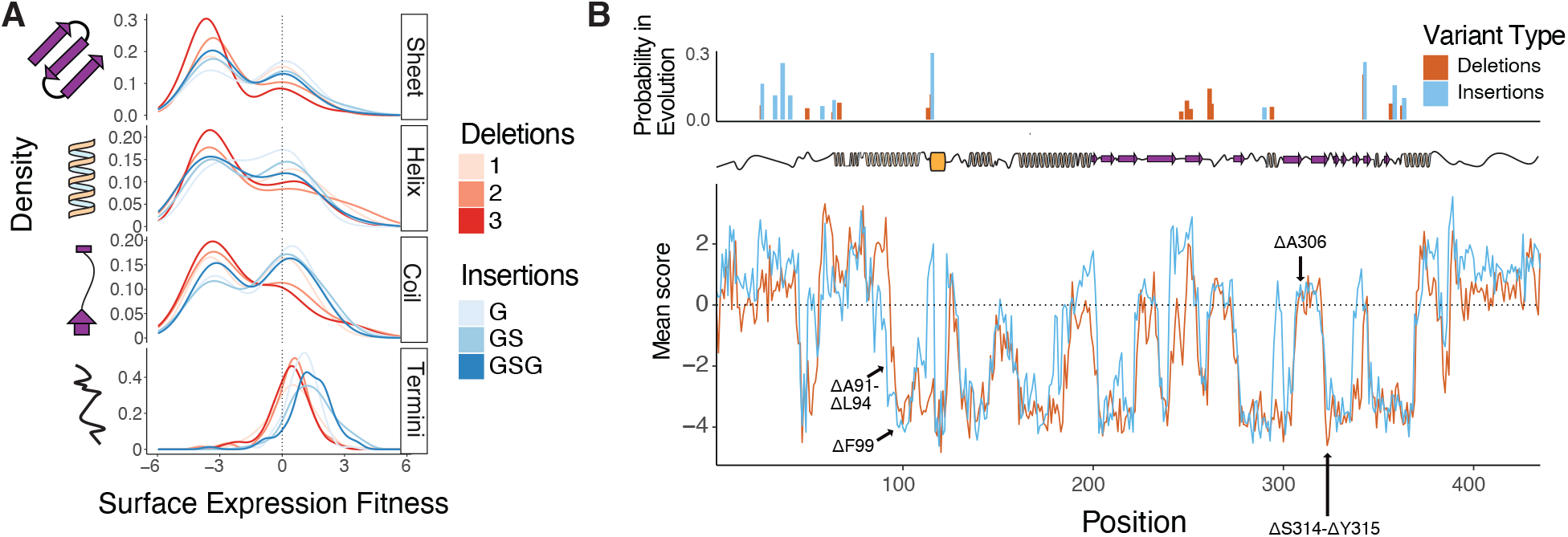
Indels have varying impact in secondary structure, disease, and evolution. **a**. The distribution of fitness effects of indels of varying length on surface expression of Kir2.1 divided by secondary structure is displayed as a kernel density estimate. Negative scores indicate decreased trafficking relative to WT Kir2.1. **b**. Mean surface scores for deletions and insertions across Kir2.1’s sequence, with conserved insertion and deletion positions in the inward rectifier protein family indicated above, in red and blue as barplots, respectively. Positions of clinically observed deletions are highlighted with arrows.

Indels occur commonly through errors in DNA replication and recombination (Kvikstad et al. 2005). As with all mutations, it’s unclear which indels are absent in extant genes due to being too deleterious or if they were never sampled in natural evolution. To compare our experimental results with natural evolution, we determined conserved indel positions in the inward rectifier family by examining the indel states of their Pfam HMM models (Mistry et al. 2021). This revealed several sites of high-probability insertion and deletion across the sequence, with slightly more deletions vs insertions (15 vs 11, Figure 6B). Given our observation that deletions are in general more deleterious than insertion, we wondered if there was a pattern to these occurrences. Many of these positions are shared, suggesting that they may be a generally permissive sites towards indels, but a distinct cluster of deletions in βE and the surrounding loops may be a site of specifically deletion-driven diversification. This agrees with our indel scanning results, which show an improved trafficking phenotype for deletions at these positions. Insertions are allowed which as well suggests that additional factors may mediate the evolutionary occurrence of indels, including mutational sampling and functional constraints. Pointing towards this, certain regions with known functional (but not trafficking) constraints, such as the CD loop, allow deletions in our data which are not observed during evolution (Coyote-Maestas et al. 2022). Conversely, a cluster of indels which are extremely deleterious in our data occur within the onset of the H helix, which suggests these positions have become specialized for Kir2.1 trafficking or folding, and perhaps harbor unknown motifs. Overall, we see insertions mostly occurring in permissive positions in our indel scan, while deletions are less tightly coupled.

## Discussion

To summarize, DIMPLE is a robust method that yields high-quality variant libraries with novel multi-codon insertions and deletions in parallel to point mutations and assayed the significance of these for trafficking of a potassium channel Kir2.1. Overall, we observe that insertions and deletions are qualitatively and quantitatively distinct from substitutions with indels more deleterious than missense mutations. By comparing indel fitness across secondary structure, we find that deletions within beta sheets are particularly deleterious while alpha helices have a range of impacts. We find that potential disease-causing deletions are highly deleterious whereas regions with allowed indels within inward rectifier diversification also allow indels in our data. Overall, our results highlight the significance of indels for mechanism, disease, and evolution.

Functional genomics approaches such as conventional deep mutational scanning and CRISPRi screens sit upon a perturbation continuum. Missense mutations inform on the physicochemical necessities of a mutated residue. In contrast CRISPRi informs you of the role of a gene within a biological network. There is a gap between the role of an individual amino acid and the entire gene such as motifs within the unstructured regions of proteins. Deletional scanning could be useful in identifying which motifs are necessary for membrane protein trafficking for instance. Indel scanning fills an important space in the functional genomics perturbation continuum.

Further work to model how indels influence proteins is clearly necessary, as existing pictures of missense mutations are not sufficient for understanding their impacts ((Holmes 2020; Tóth-Petróczy and Tawfik 2013; Savino, Desmet, and Franceus 2022)). Inserting a sequence might be akin to changing the tension on a spring, with the important parameters being the length and elasticity of the spring. Deletions have the added difficulty of removing sequences entirely, which specifically alters the registry of secondary structural elements and removes interacting residues in addition to increasing the tension on the polymer chain. We suspect an expanded framework will be necessary, considering the physical dynamics of the polymer chain itself in addition to local physicochemical changes as with substitutions. Observational bioinformatic studies of indels across evolution, have observed general trends of beta sheets having few indels, alpha helices slightly more, and flexible loops and unstructured regions having many indels (Holmes 2020; Tóth-Petróczy and Tawfik 2013; Savino, Desmet, and Franceus 2022). As with other less systematic yet still informative indel scanning studies in structured proteins, we confirm these trends. In contrast to previous indel studies in multi-domain proteins, the truly systematic nature of the data will lend itself for empirically determined models of how indels alter a protein’s backbone. Such models would be tremendously useful in understanding the fundamentals of how proteins evolve and how to engineer new proteins.

Computationally and experimental biologists are working to identify how all genetic variants impact the function of disease genes. Computationally many of the models for predicting pathogenicity use column-based multiple sequence alignments which typically do not include the gaps that indels cause. Similarly, the mutational scans which are being used for functionally characterizing variants, are mostly focused on missense mutations. Overall, this means the impact of indels are undersampled within the ongoing atlas of variant effect. We anticipate that DIMPLE will play a crucial role in filling this gap to enable the field of mutational scanning to experimentally determine how indels cause disease.

## Methods

### In silico library generation

The DIMPLE software was adapted from SPINE (Nedrud, Coyote-Maestas, and Schmidt 2021) by improving workflow and adding new functions for scanning mutations, insertions, and deletions. Additionally, for ease of use, we added a graphical user interface for those not experienced with command line interface. The first change to the code was incorporating scanning missense mutations which was adapted from a function written for a deep mutational scan (DMS) of a PDZ domain (Nedrud, Coyote-Maestas, and Schmidt 2021). We improved upon this method by adding the ability to not only mutate each position to the other 19 amino acids, but also added the option to mutate to a synonymous codon and a stop codon. These improvements are important for normalization and range in enrichment scores. The other major improvement was to add insertions and deletions at each position. Insertions are defined by the user at the nucleotide level and deletions are defined by the user as the number of nucleotides to delete. Insertions are placed following each amino acid, while deletions delete each amino acid (not including the start codon) and the next consecutive amino acids according to the length specified by the user. Therefore, the deletions stop short of the last amino acid based on the maximum length of deletions. With the addition of insertions and deletions, the size of the oligo changes and therefore needs to be buffered with additional nucleotides to match length for synthesis and for uniform amplification. Additional barcodes were used for buffering the oligo between the primer binding and the type II restriction enzyme recognition site. The size of the buffered region matched the shortest fragment (either largest deletion or smallest insertion) and was uniformly added on the 5’ and 3’ ends. Buffering at this position, however, would disrupt primer binding when using the previous SPINE software since the primer binding sites on the oligo bound partly to the type II restriction enzyme recognition site to maximize the gene fragment size. To remedy this potential issue, the barcode region was expanded so the entire primer could bind. The other changes that were made included fixing the issue of low mutation frequencies at the boundaries of the gene fragments during library generation. To generate more uniform libraries, we added overlap to each fragment by shifting the restriction sites four bases in both directions but did not add mutations in these overlaps to avoid duplication of mutations between fragments. We also added the ability to choose custom codon usage frequencies and fixed an issue with inverse PCR amplification by increasing the melting temperature threshold.

Code deposited at: https://github.com/coywil26/DIMPLE.

All primers designed and used within this manuscript for generating libraries are listed in Supplemental Table 3.

### Library generation and cloning

A SurePrint Oligonucleotide library (Agilent Technologies) containing the 58300 oligos for target genes VatD, TRPV1, and OPRM1 was synthesized by Agilent and received as 10 pmol of lyophilized DNA (Supplemental Data 2). This DNA was resuspended in 500 µL 1x TE. Sublibraries were PCR amplified using primer-specific barcodes for each sublibraries and PrimeStar GXL DNA polymerase (Takara Bio) according to the manufacturer’s instructions in 50 µL reactions using 1µL of the total OLS library as template and 25 cycles of PCR. The reactions were cleaned up using Clean and Concentrate kits (Zymo Research) and eluted in 10 μl of TE buffer. Successful amplification was assessed by running a small amount of the PCR product on an agarose gel.

Vectors containing each gene of interest were synthesized by Twist Bioscience and received as lyophilized plasmid DNA in their High Copy Number Kanamycin backbone and resuspended to 10ng/µL in 1x TE buffer. For Kir2.1 we used the same sequence we had previously used for library generation. For VatD, we designed the library with HindIII and BamHI restriction cut sites for swapping into an expression vector. For OPRM1 and TRPV1, we started with Human and Rat cDNA versions, removed BsmBI and BsaI cutsites using synonymous mutations and added flanking BsmBI cutsites which cut within CATG and GGGT on the N and C termini of each gene, respectively. These sequences were chosen so that on the N terminus of the gene we encoded for the beginning of the kozak-start codon and on the C terminus a GS linker.

For each sublibrary, the plasmid was amplified to add on golden gate compatible Type IIS restriction sites complementary to those encoded within the sublibrary oligos using Primestar GXL polymerase according to the manufacturer’s instructions in 50µL reactions using 1µL of the template vector and 25 cycles of PCR. The entire PCR reaction was run on a 0.5% agarose gel and gel purified using a Zymoclean Gel DNA Recovery Kit.

Target gene backbone PCR product and the corresponding oligo sublibrary were assembled using BsaI-mediated Golden Gate cloning. Each 40µL reaction was composed of 300ng of backbone DNA, 50ng of oligo sublibrary DNA, 2µL BsaI-HF v2 Golden Gate enzyme mixture (New England Biolabs), 4µL 10x T4 Ligase buffer, and brought up to a total volume of 40uL with nuclease free water. These reactions were placed in a thermocycler with the following program: (i) 5 min at 37°C, (ii) 5min at 16°C, (iii) repeat (i) and (ii) 29 times, (iv) 5 minutes at 60°C, (v) hold at 10°C. Reactions were cleaned using Zymo Clean and Concentrate kits, eluted into 10µL NFH2O, and transformed into MegaX DH10B (Thermo Fisher) according to manufacturer’s instructions.

Cells were recovered for one hour at 37°C. A small subset of the transformed cells were plated at varying cell density to assess transformation efficiency. All transformations had at least 100x the number of transformed colonies compared to the library size. The remaining cell outgrowth was added to 30mL LB with 50ug/mL kanamycin and grown at 37°C with shaking until the OD reached 0.6. Library DNA was isolated by miniprep (Zymo Research). Sublibrary concentration was assessed using Qubit. Each sub-library of a given gene was pooled together at an equimolar ratio. These mixed libraries were assembled with a landing pad cell line compatible backbone containing a Carbenicillin resistance cassette and GSGSGS-P2A-Puromycin cassette for positive selection.

### Sequencing library preparation and genomic DNA extraction and data analysis

Genomic DNA was extracted from sorted cells using a Micro kit from Zymo. Following DNA extraction and quantification with NanoDrop, 1.5 µg of each library was used as template for PCR using cell_line_for_3 and P2A_cell_line_rev primers with PrimeSTAR GXL enzyme, with a final primer concentration of 0.25 µM each, and a T_m_ of 56°C and 18 cycles. The amplified bands were then run on a 1.5% gel and extracted. The eluted bands were quantified using Qubit with HS kit. For VatD, samples were amplified directly from the miniprepped plasmid library using pGDP3_seq_F and pGDP3_seq_R primers, with an otherwise identical process. For OPRM1 and TrpV1 samples were amplified directly from the miniprepped plasmid library using Landing_pad_backbone_for and P2A_cell_line_rev, using the same methods.

Amplicons were prepared for sequencing using the Nextera XT DNA Library kit from Illumina with 1 ng of DNA input. Samples were indexed using the IDT for Illumina UD indexes and SPRISelect beads at a 0.9x ratio were used for cleanup and final size selection. Each indexed tagmented library was quantified with Qubit HS as well as Agilent 2200 TapeStation. Samples were then pooled and sequenced on a NovaSeq 6000 SP300 flowcell in paired-end mode, generating fastq files for each sample after demultiplexing. Each fastq was then processed in parallel using the following workflow: adapter sequences and contaminants were removed using BBDuk, then paired reads were error corrected with BBMerge and then mapped to the reference sequence using BBMap with 15-mers (all from BBTools (Bushnell 2014)). Variants in the mapped SAM file were called using the AnalyzeSaturationMutagenesis tool in GATK v4 (Van der Auwera and O’Connor 2020). The output of this tool is a csv containing the genotype of each distinct variant as well as the total number of reads. This was then further processed using a python script, which filtered out sequences that were not part of the designed variants, then formatted input files for Enrich2 (Rubin et al. 2017). Enrichment scores were calculated from the collected processed files using weighted least squares and normalized using wild-type sequences. The final scores were then processed and plotted using R. Read counts are reported within Supplemental Table 4 and Enrich2 outputs are in Supplemental Data 2.

Due to the length of synthesized oligos, microarray-based oligo library synthesis (OLS) pools typically have many errors, consisting primarily of single- and multi-base deletions (Kosuri et al. 2010; Lubock et al. 2017). Analysis of our sequencing results is consistent with this, with most off-target variants observed consisting of large deletions or frameshifts, followed by mismatches (Supplemental Figure 11, Supplemental Table 5). We observed a consistent trend where assembled products with a truncated mutagenic sublibrary were generated, with an enrichment towards the oligo beginning for larger deletions which makes sense because the oligo is synthesized from 5’-3’ ends. In previous libraries we observed an error-free final portion of ∼15%. In this work, we took advantage of an improved HiFi OLS platform from Agilent, which led to reduced error rates such that 80% of our final Kir2.1 variants consist only of our designed mutations.

The crystal structure of the closely related Kir2.2 was used to model the Kir2.1 structure (PDB: 3SPI). Homologous positions in a sequence alignment were used to map the corresponding position in the Kir2.1 sequence to the structure. An AlphaFold model of mouse Kir2.1 was examined and found to correspond closely to this method but was not used.

For the evolutionary conservation analysis, the central and C-terminal Pfam HMMs (PF01007, PF17655) were downloaded and aligned to Kir2.1. The insertion (or deletion) probability was defined as the probability of transition from a matching to an insertion (or deletion) state at each position in the profile.

### Cell line generation and cell culture

Cell lines were generated as in (Coyote-Maestas et al. 2022; Matreyek et al. 2020): prior to transfection, libraries were cloned into a landing pad vector containing a BxB1-compatible a*ttB* recombination site using BsmBI mediated golden gate cloning. We kept track of transformation efficiency to maintain library diversity that was at least 100x the size of a given library. We designed the landing pad vector which we recombined the library into to contain BsmBI cutsites with compatible overhangs for the library to have an N terminal kozak sequence and in-frame with a C-terminal GSGSGS linker-P2A-Puromycin resistance cassette. The golden gate protocol we used was 42°C for 5 minutes then 16°C for 10 minutes repeated for 35 cycles followed by 42°C for 30 minutes then 60°C for 5 minutes before being stored at 4°C prior to transfection. This landing pad backbone was generated using Q5 site-directed mutagenesis, according to the manufacturer’s suggestions.

To make the cell lines, 1000ng of library landing pad constructs were co-transfected with 1000ng of a BxB1 expression construct (pCAG-NLS-BxB1) using 3.75µL of lipofectamine 3000 and 5µL P3000 reagent in 6 wells of a 6 well plate. All cells were cultured in 1X DMEM, 10% FBS, 1% sodium pyruvate, and 1% penicillin/streptomycin (D10). The HEK293T based cell line has a tetracycline induction cassette upstream of a BxB1 recombination site and split rapamycin analog inducible dimerizable Casp-9. Two days following transfection, expression of integrated genes or iCasp-9 selection system is induced by the addition of doxycycline (2ug/µL, Sigma-Aldrich) to D10 media. Two days after induction with doxycycline, AP1903 is added (10nM, MedChemExpress) to cause dimerization of Casp9. Successful recombination shifts iCasp-9 out of frame, so only non-recombined cells will die from iCasp-9 induced apoptosis following the addition of AP1903. After two days of AP1903-Casp9 selection the media is changed back to D10 with doxycycline and cells are allowed to recover for two days.

Due to the frequent frameshifts or premature stops within OLS-based libraries we are worried they will introduce noise in our assays. To mitigate this, we typically select for proper in-frame full-length assembly by co-translationally expressing a resistance marker or fluorescent protein downstream of the target gene. This allows facile selection for variants of interest during growth or sorting. In this case we used Puromycin selection. After allowing cells to recover for two days, media was changed to D10 with doxycycline and puromycin (2ug/ml, Life Technologies Corporation), as an additional selection step to remove non-recombined cells. Cells remained in D10 plus doxycycline and puromycin for at least two days until cells stopped dying. Following puromycin treatment cells are detached, mixed, and seeded on a 10cm dish. Cells were then allowed to grow until they reached near confluence, then frozen in aliquots in a cryoprotectant media (90% FBS and 10% DMSO).

### Fluorescence-activated cell sorting

Thawed stocks of library cell lines were seeded on a 10cm dish in D10 media. The following day, the media was exchanged for fresh D10 to remove cryoprotectant media. Two days prior to the experiment, media was changed to D10 with doxycycline. After two days of induction, cells were detached with 1ml TrypLE Express (Thermo Fisher Scientific), pelleted, and washed three times with FACS buffer (5% FBS and 0.1% sodium azide in PBS). Cells were then resuspended in FACS buffer and incubated with a BV-421 anti-DYDDDDK epitope tag antibody (BioLegend) for 1 hr at 4°C. Following incubation with antibody, cells were washed two additional times with FACS buffer before being resuspended at 5 million cells per ml, filtered with cell strainer 5ml tubes (Falcon), covered with aluminum foil, and kept on ice before sorting.

All cell sorting was performed on a BD FACSAria II cell sorter. BV-421 fluorescence was excited with a 405nm laser and recorded with a 450/50 nm band pass filter. Cells were gated on forward scattering area and side scattering area to separate HEK293T whole cells then forward scattering width and height to find single cells. Surface expressed cells were separated into four subpopulations based on BV-421 fluorescence from the Anti-DYDDDDK antibody. As the library had a clear bimodal distribution, we separated up the gates based on the distribution shapes, such that the first and second gates were of the bottom and upper half of the lower fluorescence populations while the third and fourth gates were the lower and upper half of the higher fluorescence population. An example gating strategy from the FACSAria Software from the day of a sort is shown in Supplementary Figure 6. Cell collected per subpopulation is reported within Supplemental Table 4.

## Acknowledgements

We are grateful for you for taking the time to read our manuscript. We are also grateful to Matthew Howard, Eric Greene, Vijay Ramani, and Margaux Pinney, the DMS crew, and the rest of the Fraser lab for helpful feedback and discussion as we developed and conducted this project and for guidance on the manuscript. We also would like to thank the hard work of those in the UCSF Flow Cytometry Core and Center for Advanced Technology without whom none of the FACS or sequencing could have been done. This work was supported by: NIH GM123159 to JSF; a HHMI Hanna Gray Fellowship, UCSF QBI Fellowship, and Laboratory for Genomics Research pilot grant to WCM; and an NIH 1F31AI157438 to DT.

## Author contributions

This study was designed by CM, DN, JSF, and WCM. DN developed the DIMPLE software package. CM, PRG, and DT wrote the in-depth bio.protocols methods guide. CM, PRG, DT, and WCM cloned the libraries and experiments presented within this paper. CM aligned the sequences and calculated enrichments. CM and WCM analyzed the data and wrote the first draft of the manuscript maintext. CM, DN, PRG, and DT, wrote the first draft of the methods section. All authors assisted with editing and finalizing the manuscript.

