## Supplemental Figures and Tables for "Deep Insertion, Deletion, and Missense Mutation Libraries for Exploring Protein Variation in Evolution, Disease, and Biology"

The following supplemental data was used in generating this manuscript.

### Supplemental Data Files

Supplemental Data 1: **DIMPLE inputs and outputs used in testing.** Folder containing complete and the coding sequences for all genes tested and two subsequent folders: one with the complete sequences that DIMPLE was run on and one with all the output primers for amplifying sublibraries, primers to amplify the backbones and the OLS sequences to order.

Supplemental Data 2: **Agilent Sure Print OLS Pool.** Names and sequences for the OLS pools that we used to build libraries for this study.

Supplemental Data 3. **Enrich2 outputs for the Kir2.1 surface expression screen.** HGVS identifier, surface fitness scores, epsilon, and standard errors for each variant used in this study.

### Supplemental Figures

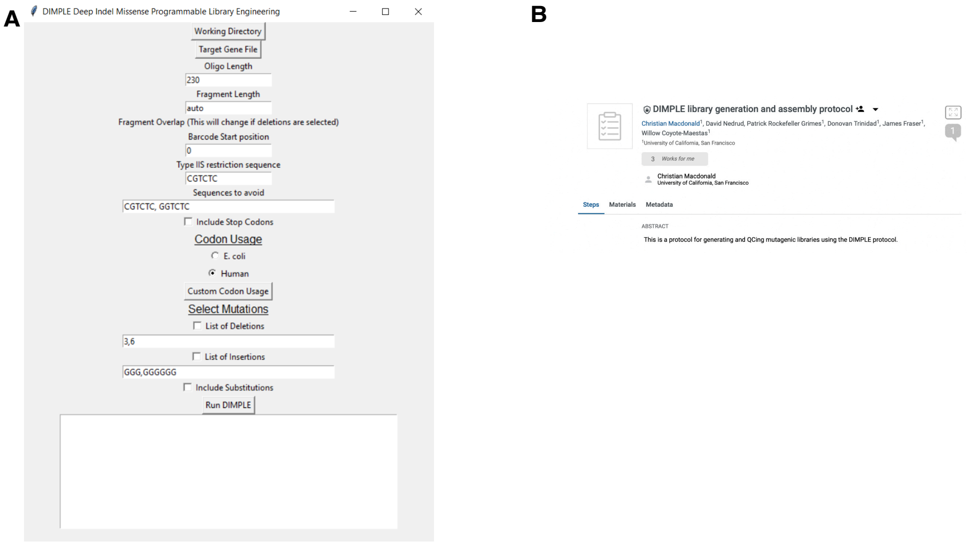

**Supplementary figure 1. DIMPLE user friendly GUI and open source protocol. A.** Open source DIMPLE graphical user interface (GUI) and detailed experimental workflow. A. We built a user friendly GUI for DIMPLE that makes library generation easy for those with little computational experience that includes many options for customization. The code is available on Github at: www.github.com/ coywil26/DIMPLE. **B.** Molecular biology can be difficult even when trying to clone a single construct. Cloning is even more challenging when trying to constructs large libraries. We've included a detailed step-by-step protocol on bioprotocols with examples that is available at: www.protocols.io/private/ 019898EFCA5611ECBEA10A58A9FEAC02.

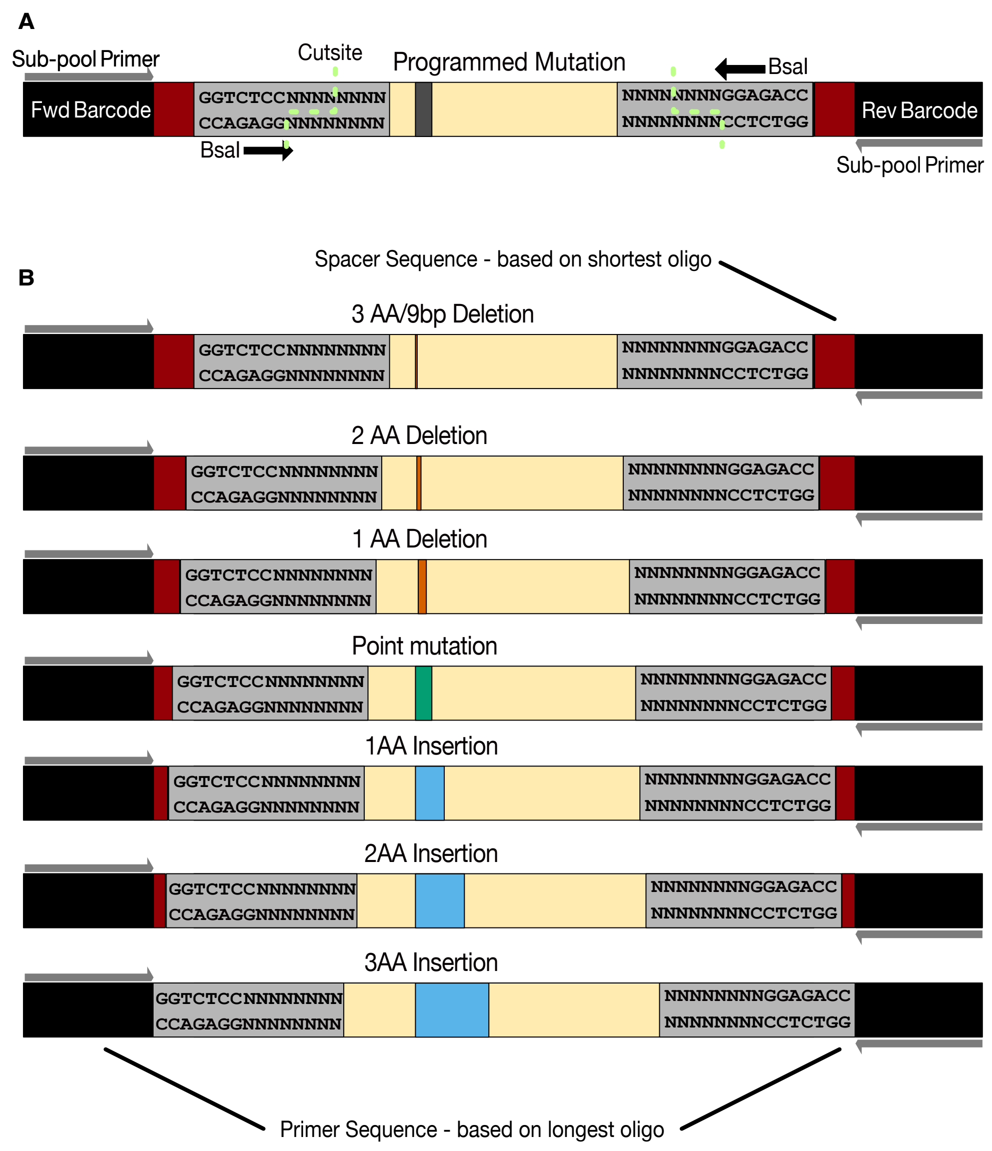

**Supplemental Figure 2. Design of oligos ordered in oligo library synthesis pools in more detail. A. Architecture of a designed mutagenic oligo.** Each sublibrary has unique forward and reverse primers for amplification of the sublibrary, spacer sequences that are adjusted to allow for the range of fragments, type IIS restriction sites, which are either BsaI or BsmBI which cut within an unmutated region of the gene of interest and having an additional 4bp of unmutated DNA beyond the cutsite to reduce bias in golden gate, and the region of a gene of interest with a single mutation. **B. Variable spacer design to reduce oligo subpool bias.** DIMPLE is designed to include deletions, insertions, and point mutations in parallel within the same subpool. The challenge is that this means there are many different sized fragments which could be amplified at different rates and could result in systematic bias. To reduce the impact of length bias from PCR, we include a variable lengthed spacer sequence (shown in red) which is defined based on the shortest oligo to bring the length equivalent to the longest oligo. To make sure that primer sequences don't change, the subpool amplifying primer sequence is determined based on the longest oligo then added to the ends of the other oligos within the subpool.

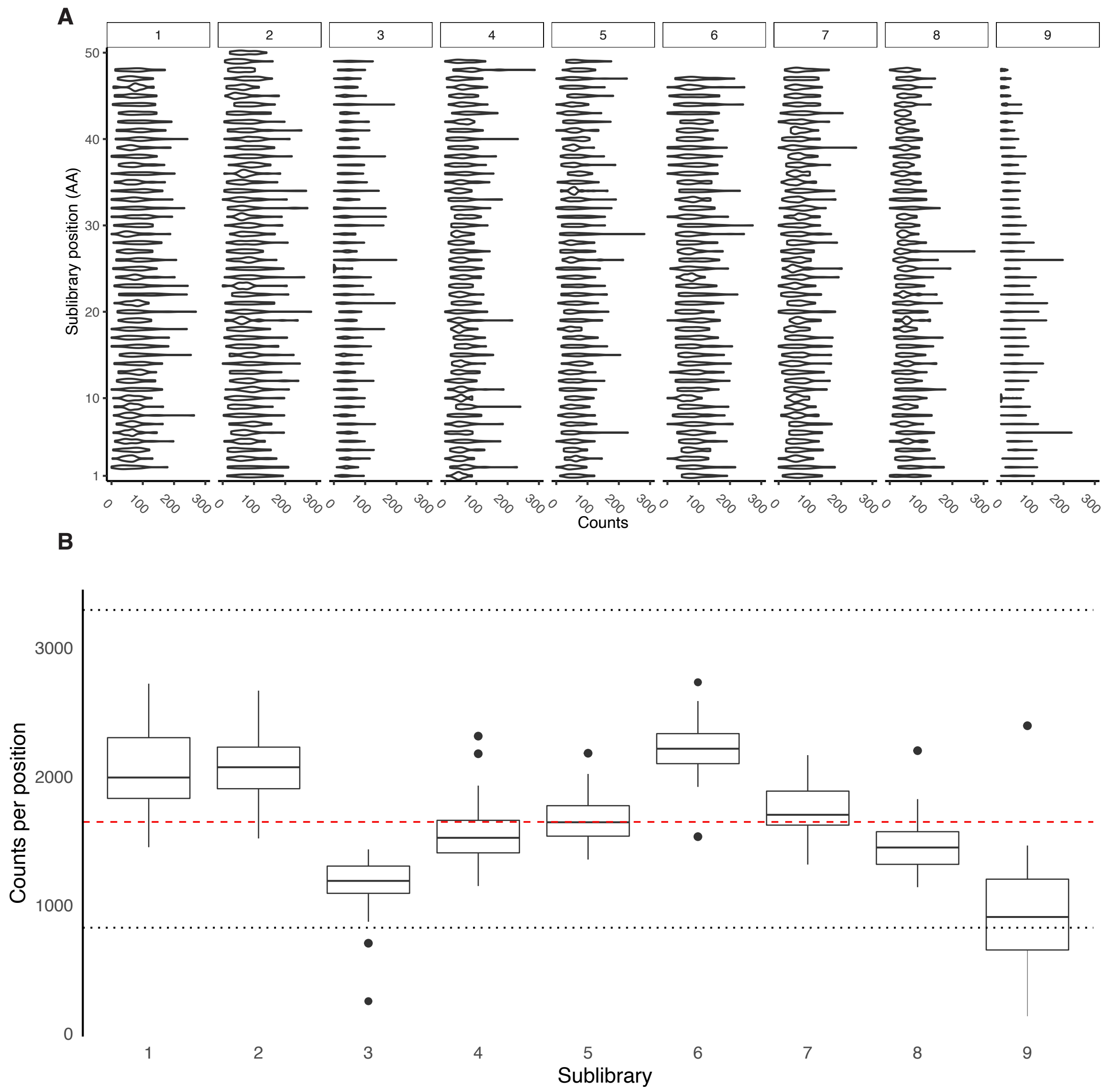

**Supplemental Figure 3. Kir2.1 Positional coverage. A.** Violin plots of every positions across all 9 sublibraries of Kir2.1. These are symmetric density plots that shows the distribution of observed read counts for all variant types at a given position. Overall, we see even representation within each sublibrary for variants, with no systematic bias for any particular region. Any variance between positions is likely more due to difficulties in oligo synthesis or sequencing. There is slightly lower read counts for the end of sublibrary 9 which likely due to tagmentation's well known bias away from the ends of a fragment of DNA. **B.** Boxplots of coverage across Kir2.1 broken apart based on sublibrary. The vertical length of the box is the interquartile range (IQR), upper bound is the 75th percentile with the lower bound is the 25th percentile. Dashed red line indicates mean overall counts per positions across subpools and dotted black lines are 2-fold and 1/2-fold the mean.

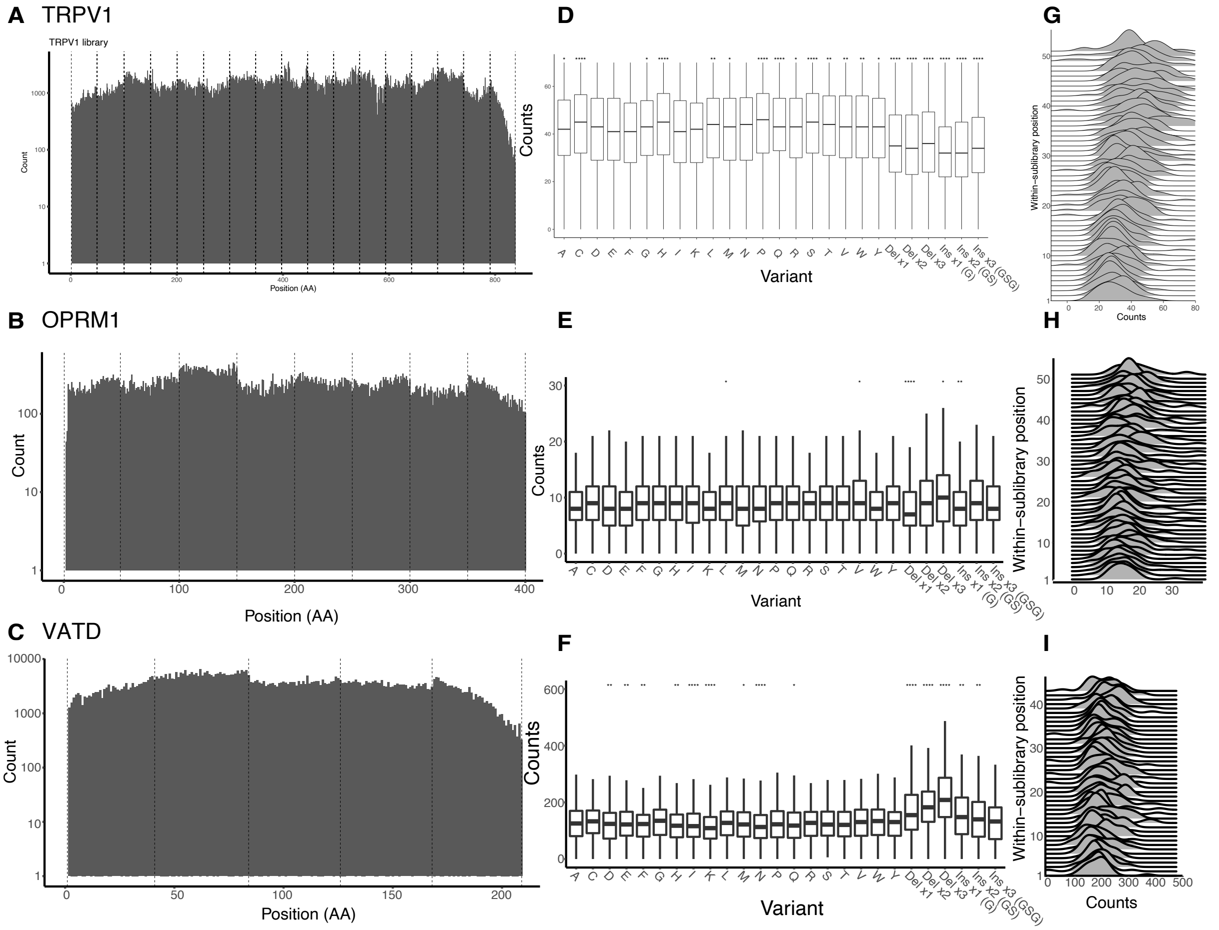

**Supplemental Figure 4. A-C.** Barplot of counts of mutation numbers per position for TrpV1, OPRM1, and VatD. The boundaries of each mutagenic sublibrary are indicated with dashed lines. Overall, each mutagenized library is pretty even across subpools, with TrpV1, OPRM1, and VatD all being within x, y, and z fold between medians for a given chunk, respectively. Sequencing libraries were constructed using Nextera which results in poor coverage at the edges which is likely particularly impacting TrpV1 sequencing as the edges of the DNA fragment were closer than ideal. **D-F.** Boxplots of variants at each position across all of TrpV1, OPRM1, and VatD. The vertical length of the box is the interquartile range (IQR), upper bound is the 75th percentile with the lower bound is the 25th percentile. Significance is tested using two-sided t-tests controlled for multiple comparisons comparing incorporation means between variants across all positions. Significance levels: ***P<0.001; **P<0.01; *P<0.05, all others not significant. Across the all libraries library, different variants are incorporated at similar rates. Sequencing-based counts of designed variant incorporation show small quantitative differences between variants, with different classes being present within x, y, z -fold of each other in TrpV1, OPRM1, and VatD libraries, respectively. Insertions and deletions appear slightly depleted compared to all other variant types in TrpV1. OPRM1 deletions appear to be slightly enriched whereas all other variant types are present at similar frequencies. VatD deletions seem to be slightly enriched as well. Overall, variant types in many different backgrounds are present at similar frequencies. **G- I.** Stacked density plots, or ridge plot ordered bottom-to-top from first to last positions of the second sublibrary of TrpV1, OPRM1, VatD. The second sublibrary was included here because the first and last sub libraries have lower sequencing coverage due to well known reduced tagementation efficiency at DNA ends. Overall, we find little positional bias at the beginnings and ends of a sublibrary, which we previously observed in OLS based libraries meaning adding a 4bp unmutated sequence appears to have reduced end based bias.

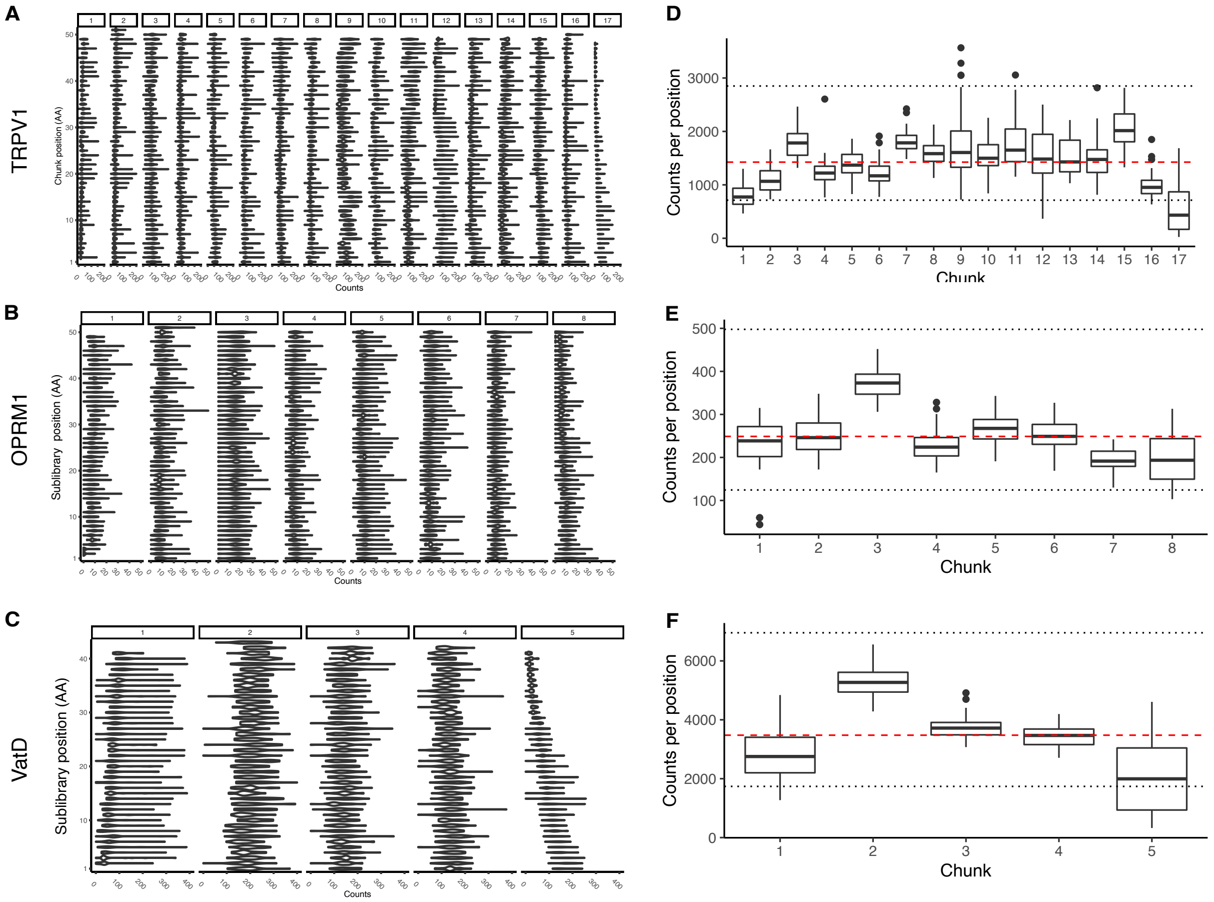

**Supplemental Figure 5. A-C.** Violin plots of every positions across all 17, 9, and 5 subpools of TrpV1, OPRM1, and VatD genes, respectively. These are symmetric density plots that shows the distribution of observed read counts for all variant types at a given position. Any variance between positions within a subpool is likely more due to difficulties in oligo synthesis or sequencing. Subpool 9 is much smaller than the others, hence why it ends after position 37. There is slightly lower read counts for the end of sublibraries 1 and 17 in TrpV1 and 5 in VatD which are likely due to Nextera's tagmentations low efficiency at the ends of a fragment of DNA. Overall, we see even representation within each subpool for variants, with no systematic bias for any particular region of across subpools. D-F**.** Boxplots of coverage across TrpV1, OPRM1, and VatD broken apart based on sublibrary. The vertical length of the box is the interquartile range (IQR), upper bound is the 75th percentile with the lower bound is the 25th percentile. Dashed red line indicates mean overall counts per positions across subpools and dotted black lines are 2-fold and 1/2-fold the mean. All libraries except for TrpV1 1 and 17 are within two fold the mean, which is likely due to tagmenetation having low efficiencies at the edges of a DNA fragment.

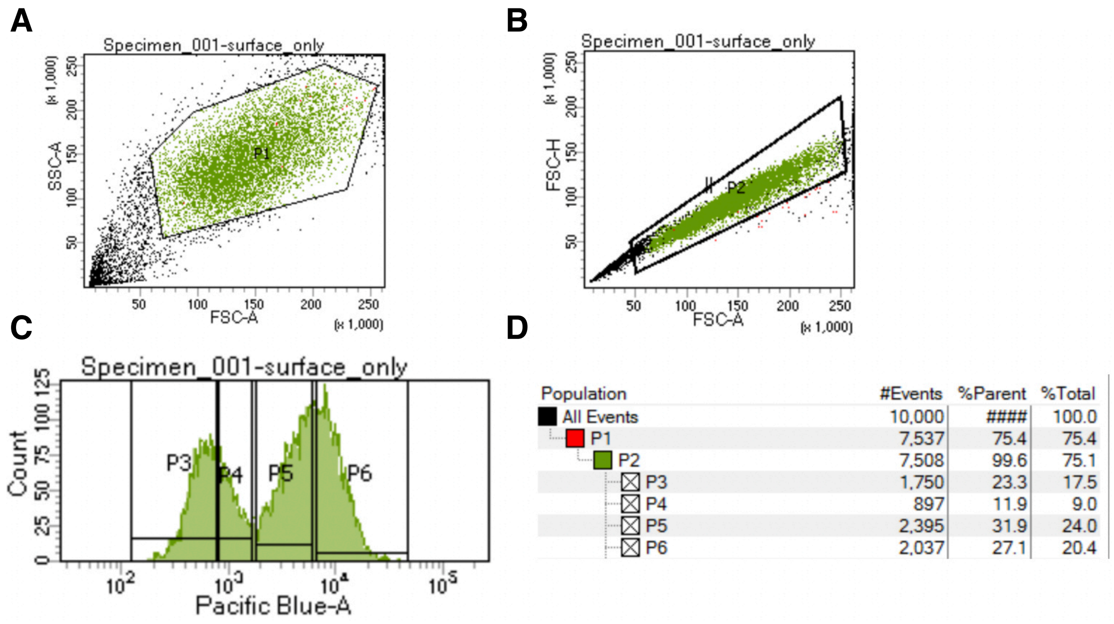

| Population | #Events | %Parent | %Total |
|---|---|---|---|
| All Events | 10,000 | #### | 100.0 |
| P1 | 7,537 | 75.4 | 75.4 |
| P2 | 7,508 | 99.6 | 75.1 |
| P3 | 1,750 | 23.3 | 17.5 |
| P4 | 897 | 11.9 | 9.0 |
| P5 | 2,395 | 31.9 | 24.0 |
| P6 | 2,037 | 27.1 | 20.4 |

**Supplemental Figure 6. Gating strategy for sorting Kir2.1 DIMPLE libraries based on surface expression. A.** Whole HEK293T cells were gated on forward and side scattering area. **B.** Single cells were gated on forward scattering area and height. CBV-421 fluorescence was excited with a 405nm laser and recorded with a 450/50nm band pass filter. **C.** Cells were then separated into four gates based on increasing BV-421 fluorescence. **D.** Cell counts from FACS experiment in sample of 10,000 events.

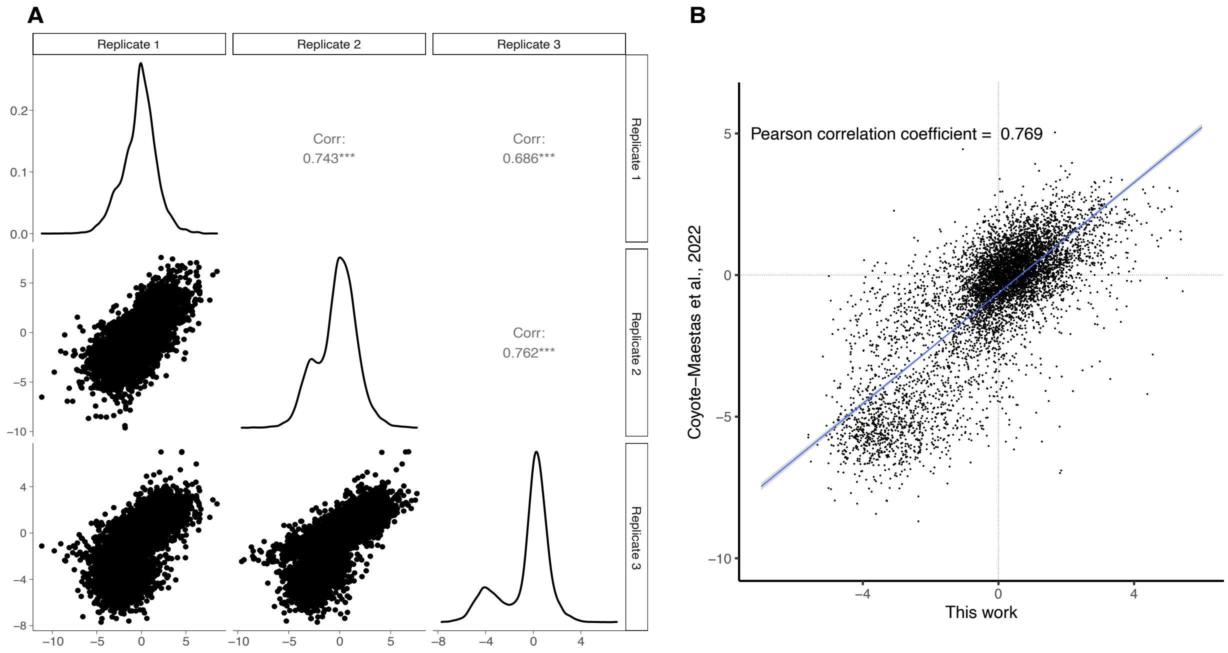

**Supplemental Figure 7. Replicates within and outside of study have high agreement. A.** Correlation plots between single replicate fitness scores and fitness distributions for all three replicates. We find good replicability with replicate 1 and 3 most different. As these experiments were done on different days over the course of a week perhaps expression levels were changing. It appears that replicate 1 is less bimodal then replicate 3 which could indicate reduced expression levels and therefore dynamic range. Overall, however we find good agreement between each biological replicate. **B.** As we previously have done a missense mutational scan of Kir2.1, we wanted to test whether we saw agreement. Indeed we see very good agreement between fitness scores. Interestingly, it appears that the overall fitness scores from the experiments are more similar based on a higher pearson correlation coefficient (0.769 between studied vs 0.762-0.686 between biological replicates). This implies that high repeatability across experiments for the Kir2.1 surface expression screen.

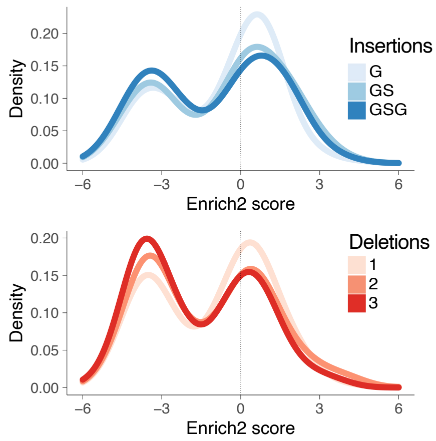

**Supplemental Figure 8. Distributions** The distribution of fitness effects on surface expression of Kir2.1 from different length insertions and deletions are displayed as kernel density estimates. Negative scores indicate decreased trafficking relative to WT Kir2.1 based on synonymous mutations. As we increase the length of an insertion or deletion, it becomes more deleterious. Overall deletions are more disruptive than insertions.

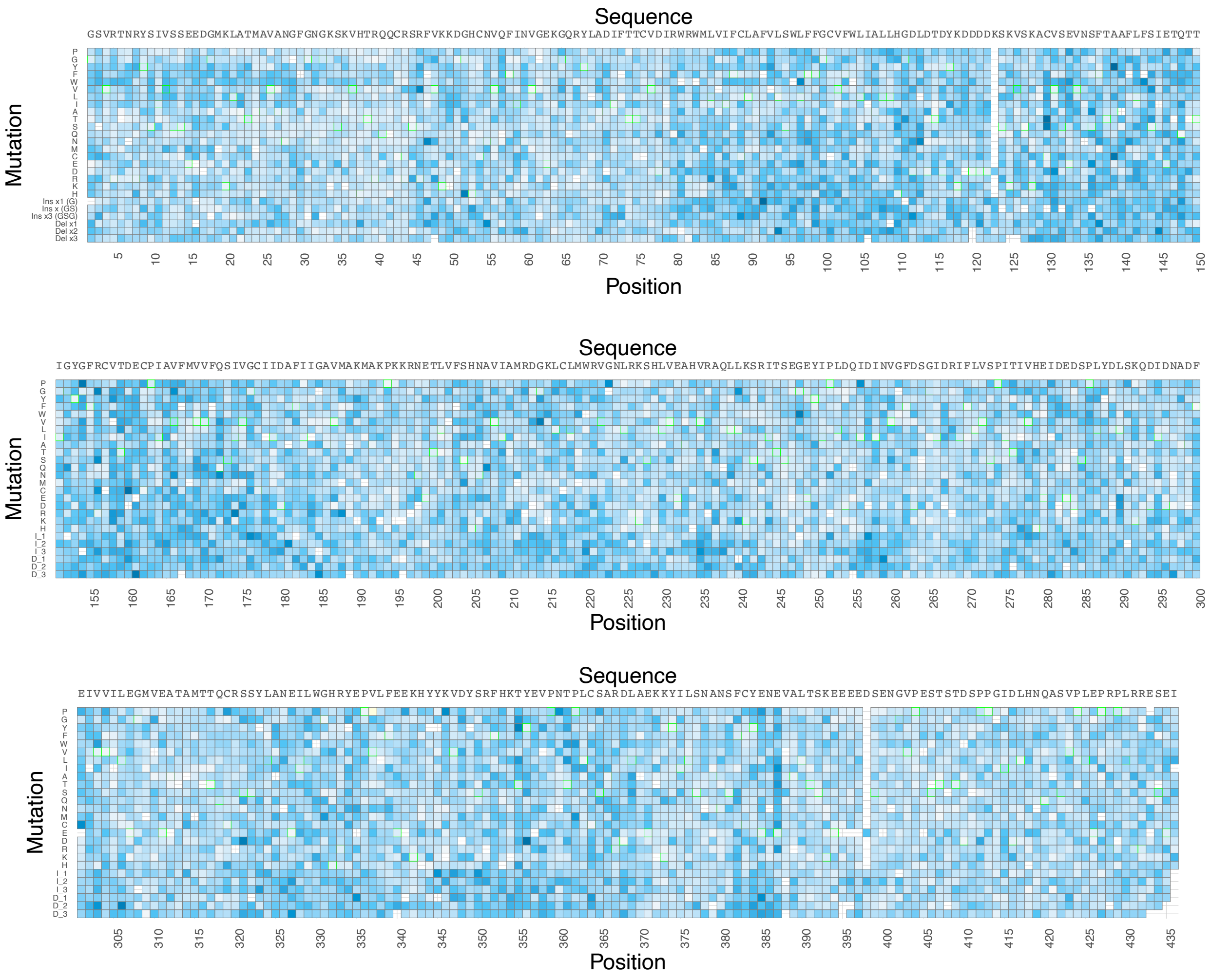

**Supplemental figure 9. Kir2.1 standard error are evenly distributed.** Heatmap of surface expression standard errors scores calculated from Enrich2 gradient colored from white-to-blue . Only positions for which there were reads in all three replicates are shown here; others were removed in enrichment calculations. Synonymous mutation boxes are outlined with green. Overall, standard error is even across the gene and relatively low meaning the data is high quality.

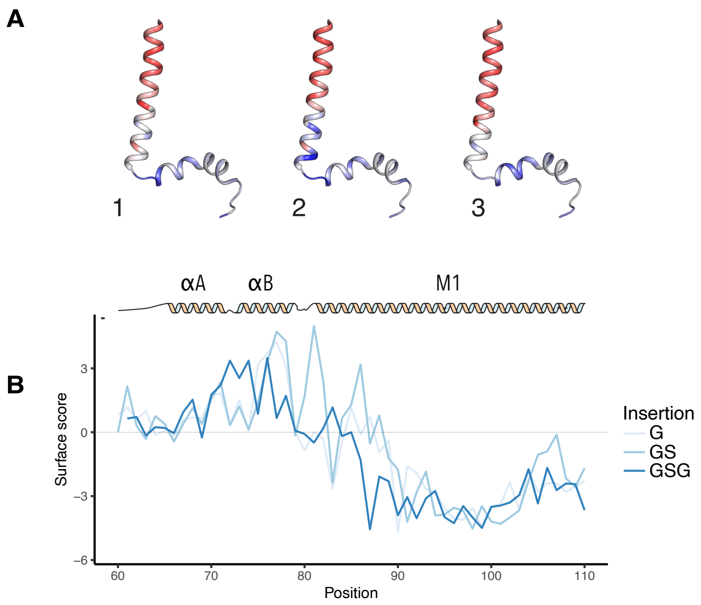

**Supplemental figure 10. Impact of varying insertion length on M1 and slide helix. A.** Impact of varying the length of insertion on surface expression mapped onto transmembrane helix M1 and slide helix colored from low- to-neutral-to-high surface expression, red-to-white-to-blue, respectively. **B.** Surface scores for the slide-helix position with varying lengths of insertion colored with increasing hue for increasing length. All insertions are poorly tolerated within the transmembrane likely due to increasing flexibility. Flexible linkers are tolerated within the extracellular loop which intuitively makes sense.

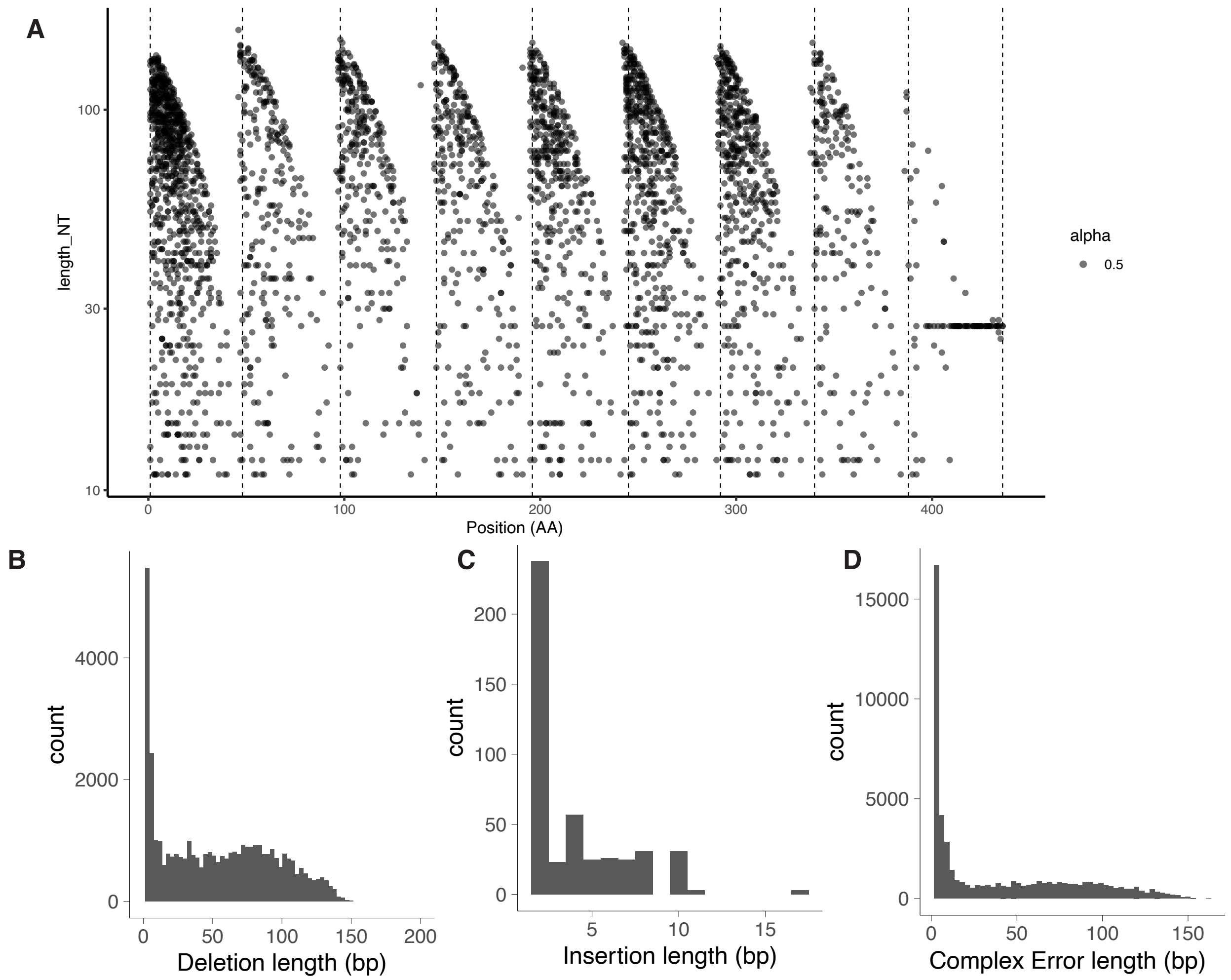

**Supplemental Figure 11. Length of deletion errors within Kir2.1 OLS subpools. A.** Plotted are the observed length of deletion variants within sublibraries that are not designed within the sublibrary against the position of the codon where it starts. The sublibraries boundaries are indicated by the dashed line. Agilent OLS pools have substantial mutational burden including many deletion variants. Because the oligos are synthesized from 5'-3' we would expect deletions to start at the 5' end and decay because it is not possible to have a deletion across sublibrary boundaries. That we see the clean decay implies errors within the libraries are due to oligo synthesis and not our cloning pipeline. Over time as technology improves so will the quality of the libraries. **B-D.** Frequency of errors based of different length non-designed deletions and insertions within the baseline Kir2.1 library. In OLS libraries single base pair deletions are the most common error type which can be clearly seen in **B**. However, frequently there are most frequently multiple errors at once within a mutated oligo that contain single base pair deletions hence the high error count in **D**.

### Supplemental Tables

| Name | Length | GC content | Variants | Sublibraries |
|---|---|---|---|---|
| ABCG2 | 1968 | 43 | 17056 | 13 |
| Amyloid beta(1-42) | 129 | 43 | 1118 | 1 |
| ASIC1a | 1638 | 48 | 14196 | 12 |
| ASIC2a | 1560 | 46 | 13520 | 12 |
| DHFR | 564 | 49 | 4888 | 4 |
| GPR161 | 1680 | 53 | 14560 | 12 |
| Hikeshi | 594 | 43 | 5148 | 5 |
| Kir2.1 | 1320 | 47 | 11440 | 9 |
| Kir2.2 | 1323 | 49 | 11466 | 10 |
| Kir3.1 | 2304 | 45 | 19968 | 11 |
| Kir3.2 | 2052 | 52 | 17784 | 9 |
| Kir6.2 | 1947 | 53 | 16874 | 9 |
| Kv1.3 | 2502 | 54 | 21684 | 13 |
| Nachra7 | 2313 | 59 | 20046 | 11 |
| OCT1 | 1743 | 57 | 15106 | 11 |
| OPRM1 | 1146 | 49 | 9932 | 8 |
| P2X3 | 1992 | 52 | 17264 | 9 |
| P2X4 | 1965 | 53 | 17030 | 9 |
| Piezo1 | 7683 | 58 | 66586 | 54 |
| Shaker | 1848 | 47 | 16016 | 13 |
| Tem1 | 861 | 52 | 7462 | 6 |
| TRPV1 | 3531 | 47 | 30602 | 17 |
| TSHR | 2322 | 48 | 20124 | 16 |
| VatD | 630 | 44 | 5460 | 5 |

Supplemental Table 1. **Inputs and outputs from test running DIMPLE.** As discussed within the text, we tested the computational DIMPLE pipeline by including 25 genes. Within this table is their length, GC content, number variants DIMPLE generated, and the number of sub libraries for the gene.

| Reagent | For 1 sublibrary | cost | total used | total |
|---|---|---|---|---|
| Zymo clean and concentrate | 1 | 2 | 9 | 18 |
| Zymo gel purification | 1 | 2 | 9 | 18 |
| MegaX competent cells | 1 | 16 | 10 | 160 |
| GXL pcr mixes | 2 | 0.75 | 9 | 13.5 |
| Sanger sequencing | 15 | 4 | 11 | 660 |
| Agilent Oligos | 1248 | 0.17 | 9 | 1909.44 |
| Zymo mini-preps | 4 | 0.91 | 10 | 36.4 |
| NEB golden gate | 10 | 4.2 | 10 | 420 |
| | total number variants | 11232 | total cost | 3235.34 |
| | | | cost per variant | 0.288046652 |

Supplemental Table 2. **Cost for generating DIMPLE libraries**. Listed are the reagents used in generating the Kir2.1 library, the number of these reagents used per sublibrary, their estimated cost based on list prices (not-institutionally agreed discounts) as of July 26 2022, the number of reactions for a 9 sublibrary gene 436 amino acids long such as Kir2.1, and total cost for that reagent across the library. Below is listed the total estimated cost for Kir2.1, $3235.34, and cost per variant, $0.28. Both are likely large over-estimated for academics due to using list prices.

| Name | Sequence |
|---|---|
| **Primers for amplifying samples** | |
| cell_line_for_3 | CCCTATCAGTGATAGAGAACGTATGACC |
| P2A_cell_line_rev | TGGTCCTGGATTTTCCTCCACATC |
| Landing_pad_backbone_for | actcactatagggcgaattggg |
| pGDP3_seq_65_F | taaccctgataaatgcttctctagaaataattttgtttaactttaaga |
| pGDP3_seq_65_R | CTTTGTTAGCAGCCGGATCTCAGTGGT |
| **Library generation Primers** | |
| TrpV1_oligoP_DMS-1_F Frag2-49_56.7C | ATATAGATGCCGTCCTAGCG |
| TrpV1_oligoP_DMS-1_R Frag2-49_56.8C | TTCCTGTGCCCATGGTAGTAAT |
| TrpV1_oligoP_DMS-2_F Frag50-100_58.6C | TTGGTCATGTGCTTTTCGGG |
| TrpV1_oligoP_DMS-2_R Frag50-100_55.9C | ACTCAGAATCGATTCCCCG |
| TrpV1_oligoP_DMS-3_F Frag101-150_57.0C | TCCGACGGGGAGTATATACGG |
| TrpV1_oligoP_DMS-3_R Frag101-150_58.2C | TCAGGCCAAAATAATCTCCG |
| TrpV1_oligoP_DMS-4_F Frag151-200_57.6C | GTACATGAAACGATGGACGGG |
| TrpV1_oligoP_DMS-4_R Frag151-200_58.8C | TCCTCAGCGATGATCCTATGG |
| TrpV1_oligoP_DMS-5_F Frag201-250_56.5C | GGCGAGAGGAGATATAGAGGG |
| TrpV1_oligoP_DMS-5_R Frag201-250_56.5C | GCTGTGCACATATATGGTCAGG |
| TrpV1_oligoP_DMS-6_F Frag251-299_56.9C | TTATAATCATCCTCCCCGGCT |
| TrpV1_oligoP_DMS-6_R Frag251-299_58.9C | TGTGTGCTCGACAGTTGACAC |
| TrpV1_oligoP_DMS-7_F Frag300-348_56.7C | AAGTTTGTGGGGTAGGGAATGG |
| TrpV1_oligoP_DMS-7_R Frag300-348_56.9C | TACCTTTGCTAGGGAGCATAAG |
| TrpV1_oligoP_DMS-8_F Frag349-397_56.5C | TGTCAGGGGTAGTGTAACACTGG |
| TrpV1_oligoP_DMS-8_R Frag349-397_58.4C | GAAATCGCTTCTAGGCAAGTTGG |
| TrpV1_oligoP_DMS-9_F Frag398-446_56.3C | AGACCAGGATGGCTGATAAGT |
| TrpV1_oligoP_DMS-9_R Frag398-446_56.1C | CACATATACCATCTACACCACGG |
| TrpV1_oligoP_DMS-10_F Frag447-495_56.9C | ATGACCGTTAGTGAGGCTAGT |
| TrpV1_oligoP_DMS-10_R Frag447-495_56.5C | TGTCCGTCTGATGGGTAATTTGG |
| TrpV1_oligoP_DMS-11_F Frag496-544_57.1C | TGGCCAGATTAATTCCCAGTTGG |
| TrpV1_oligoP_DMS-11_R Frag496-544_56.0C | ATCTTTATGGTTCTCCCACATGG |
| TrpV1_oligoP_DMS-12_F Frag545-593_56.6C | TCTCATTGTCTATTCCCCGGT |
| TrpV1_oligoP_DMS-12_R Frag545-593_58.2C | CTCAAGTACCGACAATCATGC |
| TrpV1_oligoP_DMS-13_F Frag594-642_58.4C | TCTTGTTGTGATGGGAGTCAAGG |
| TrpV1_oligoP_DMS-13_R Frag594-642_56.5C | GTCGTCAGCCGCCTATG |
| TrpV1_oligoP_DMS-14_F Frag643-691_56.8C | AATGGTACATCCCTAGCAGGAG |
| TrpV1_oligoP_DMS-14_R Frag643-691_57.1C | TCGATGCCGTTTTTGACCC |
| TrpV1_oligoP_DMS-15_F Frag692-740_57.5C | TACAGGGATTTCGAGGCTTTAG |
| TrpV1_oligoP_DMS-15_R Frag692-740_58.7C | GCTCGGTCTGTATCTCTTAACG |
| TrpV1_oligoP_DMS-16_F Frag741-789_59.0C | CTCAGCTGCTCACTGGTATATGG |
| TrpV1_oligoP_DMS-16_R Frag741-789_57.7C | GTCTGAAAGTCCGGGTGATCT |
| TrpV1_oligoP_DMS-17_F Frag790-838_57.9C | GAACTTGATGTGGCAGTCTGTGG |
| TrpV1_oligoP_DMS-17_R Frag790-838_56.6C | GGGTCCCTAGAAAAGTCATCAG |
| TrpV1_geneP_Mut-1_R Frag2-49 58.1C | ATAGGTCTCGTGAGACGGCcggatctg |

| | |
|---|---|
| TrpV1_geneP_Mut-1_F Frag2-49 59.0C | ATAGGTCTCGTAAGGGCGATAGCGAGGA |
| TrpV1_geneP_Mut-2_R Frag50-100 56.8C | ATAGGTCTCCTAGTCCTGGACCGTGTG |
| TrpV1_geneP_Mut-2_F Frag50-100 59.1C | ATAGGTCTCTTTCTGCCGGGGAAAAGCC |
| TrpV1_geneP_Mut-3_R Frag101-150 60.3C | ATAGGTCTCTGGGAAGAAGGTCTAACACTAGC |
| TrpV1_geneP_Mut-3_F Frag101-150 59.2C | ATAGGTCTCAGACTGGCAAAACCTGTTTGCT |
| TrpV1_geneP_Mut-4_R Frag151-200 59.0C | ATAGGTCTCTTGAACTCACTGTCGGTCAACC |
| TrpV1_geneP_Mut-4_F Frag151-200 56.3C | ATAGGTCTCAAACCGCCCTCCATATCG |
| TrpV1_geneP_Mut-5_R Frag201-250 60.4C | ATAGGTCTCTAATAAGAGTCGGTGTAGCTGGC |
| TrpV1_geneP_Mut-5_F Frag201-250 59.6C | ATAGGTCTCTTTCCCTGGCTGCATGC |
| TrpV1_geneP_Mut-6_R Frag251-299 56.5C | ATAGGTCTCTCCCCGAAGTAGAATCCAGG |
| TrpV1_geneP_Mut-6_F Frag251-299 60.1C | ATAGGTCTCACACTAAGTTCGTTACCAGTATGTATAAC |
| TrpV1_geneP_Mut-7_R Frag300-348 59.2C | ATAGGTCTCGTATTGTCAGCCACTTCGACCA |
| TrpV1_geneP_Mut-7_F Frag300-348 61.1C | ATAGGTCTCCTTATATATTGCAACGAGAGATACATGAGC |
| TrpV1_geneP_Mut-8_R Frag349-397 59.1C | ATAGGTCTCCCTATTTTTCCAGAACTGGCAGC |
| TrpV1_geneP_Mut-8_F Frag349-397 62.0C | ATAGGTCTCTCGCTTACTCAAGTAGCGAAACC |
| TrpV1_geneP_Mut-9_R Frag398-446 59.3C | ATAGGTCTCAGGACGCTGTTTTTCTCACATGT |
| TrpV1_geneP_Mut-9_F Frag398-446 60.2C | ATAGGTCTCTCACCGCAGCAGCTTATTACC |
| TrpV1_geneP_Mut-10_R Frag447-495 59.1C | ATAGGTCTCATATACAGGCAATACACAAAGAAATTGA |
| TrpV1_geneP_Mut-10_F Frag447-495 60.2C | ATAGGTCTCTGCAGCGCAGACCTTCTTT |
| TrpV1_geneP_Mut-11_R Frag496-544 60.3C | ATAGGTCTCTGGATGCCTCTAAAAAAGAAGTACAC |
| TrpV1_geneP_Mut-11_F Frag496-544 55.7C | ATAGGTCTCCAATGGGATGGACCAATATGTTGT |
| TrpV1_geneP_Mut-12_R Frag545-593 60.4C | ATAGGTCTCAAAACCATAGAAGCAACGTACTCTTTG |
| TrpV1_geneP_Mut-12_F Frag545-593 59.1C | ATAGGTCTCTCGTAACACTTATAGAGGACGGG |
| TrpV1_geneP_Mut-13_R Frag594-642 60.5C | ATAGGTCTCGAGAATCCGAAGAGGAAAACAAGGT |
| TrpV1_geneP_Mut-13_F Frag594-642 57.9C | ATAGGTCTCTGGGCGACCTTGAGTTCAC |
| TrpV1_geneP_Mut-14_R Frag643-691 60.0C | ATAGGTCTCGTGAATTTAAAGAGTTCCAGGCATGTG |
| TrpV1_geneP_Mut-14_F Frag643-691 60.3C | ATAGGTCTCGCAAGAACATTTGGAAGCTCCAAAG |
| TrpV1_geneP_Mut-15_R Frag692-740 61.7C | ATAGGTCTCGCGATTTTATTGACGGTTTCGCC |
| TrpV1_geneP_Mut-15_F Frag692-740 58.9C | ATAGGTCTCTTAGGGTAGACGAAGTTAACTGGA |
| TrpV1_geneP_Mut-16_R Frag741-789 56.6C | ATAGGTCTCCGATAATCATCCTTCCCGTC |
| TrpV1_geneP_Mut-16_F Frag741-789 56.0C | ATAGGTCTCCCCTCGTGCCACTCCTT |
| TrpV1_geneP_Mut-17_R Frag790-838 57.8C | ATAGGTCTCTTCCAGTTGCGGCCACTG |
| TrpV1_geneP_Mut-17_F Frag790-838 61.7C | ATAGGTCTCCGAGACGtGCAACTAATTTTAGTCTAC |
| Kir21_oligoP_DMS-1_F Frag2-48_56.2C | AAGACTCAACCAATGACCCTTC |
| Kir21_oligoP_DMS-1_R Frag2-48_55.6C | CAATACTGAGCCGTGGGTT |
| Kir21_oligoP_DMS-2_F Frag49-98_56.0C | ACAATAACTATGGGTGCGGGG |
| Kir21_oligoP_DMS-2_R Frag49-98_57.9C | CCCGATGGATCTTTATACTCGG |
| Kir21_oligoP_DMS-3_F Frag99-147_57.4C | TGGTTGTATAGCTGAGCGATG |
| Kir21_oligoP_DMS-3_R Frag99-147_59.0C | GAGAGTTGCGCCGAGTTTG |
| Kir21_oligoP_DMS-4_F Frag148-196_57.9C | AGCTAAGACAATACCCTCGGAGG |
| Kir21_oligoP_DMS-4_R Frag148-196_57.2C | CCGTAAGCTCTAGAGGTTGTC |
| Kir21_oligoP_DMS-5_F Frag197-244_56.1C | TGGTGATAGGTAAGGATGGCA |
| Kir21_oligoP_DMS-5_R Frag197-244_58.0C | CATAATATCTGACGTTAGGCGTTGG |
| Kir21_oligoP_DMS-6_F Frag245-292_56.6C | GTCGATGGCTCCCCTTATTAC |

| | |
|---|---|
| Kir21_oligoP_DMS-6_R Frag245-292_59.3C | GACAGTCTCTCGTTTGTTCCTTAGG |
| Kir21_oligoP_DMS-7_F Frag293-340_56.4C | AGTTAGTGGTTGCCTCAGAATC |
| Kir21_oligoP_DMS-7_R Frag293-340_57.9C | ATCGCATCCCATTATCTAGCG |
| Kir21_oligoP_DMS-8_F Frag341-388_55.7C | GCCTTTATGTTCCATGGTCCT |
| Kir21_oligoP_DMS-8_R Frag341-388_58.5C | CGTCTAGCCCAGAGAGAGTC |
| Kir21_oligoP_DMS-9_F Frag389-436_58.6C | GAGCGGATCTGGATACGTAATTGG |
| Kir21_oligoP_DMS-9_R Frag389-436_57.7C | TTAGACTTGCAGTCTCATAATGGG |
| Kir21_geneP_Mut-1_R Frag2-48 58.1C | ATAGGTCTCGTGAGACGGCcggatctg |
| Kir21_geneP_Mut-1_F Frag2-48 57.8C | ATAGGTCTCAAGATGGCCACTGTAATGTGCA |
| Kir21_geneP_Mut-2_R Frag49-98 58.1C | ATAGGTCTCAAGCGTGACCGGCACTG |
| Kir21_geneP_Mut-2_F Frag49-98 61.7C | ATAGGTCTCGCTGCGTTTTTTGGTTGATCGC |
| Kir21_geneP_Mut-3_R Frag99-147 59.5C | ATAGGTCTCAGCCAAGACAGTACGAAGGC |
| Kir21_geneP_Mut-3_F Frag99-147 56.2C | ATAGGTCTCCCACTATCGGATACGGGTTC |
| Kir21_geneP_Mut-4_R Frag148-196 61.7C | ATAGGTCTCTCGATAGAGAAAAGGAAAGCTGCG |
| Kir21_geneP_Mut-4_F Frag148-196 60.5C | ATAGGTCTCATGAGACATTGGTTTTCAGTCACAACG |
| Kir21_geneP_Mut-5_R Frag197-244 58.3C | ATAGGTCTCTTGGGCTTAGCCATTTTTGCC |
| Kir21_geneP_Mut-5_F Frag197-244 59.4C | ATAGGTCTCCAGAAGGAGAGTACATACCACTCG |
| Kir21_geneP_Mut-6_R Frag245-292 61.7C | ATAGGTCTCCGTGATTTCAGCAGTTGTGCTCG |
| Kir21_geneP_Mut-6_F Frag245-292 60.5C | ATAGGTCTCACATCGACAACGCTGATTTCGAA |
| Kir21_geneP_Mut-7_R Frag293-340 60.2C | ATAGGTCTCGAGAGGTCATAAAGAGGAGAATCCTC |
| Kir21_geneP_Mut-7_F Frag293-340 61.1C | ATAGGTCTCAGCACTATTATAAGGTCGACTACTCTAG |
| Kir21_geneP_Mut-8_R Frag341-388 60.4C | ATAGGTCTCAACAAGACAGGTTCGTAGCGATG |
| Kir21_geneP_Mut-8_F Frag341-388 61.2C | ATAGGTCTCTCACATCAAAGGAAGAAGAAGAAGATT |
| Kir21_geneP_Mut-9_R Frag389-436 62.7C | ATAGGTCTCTCGTTCTCGTAACAGAAAGAGTTTGC |
| Kir21_geneP_Mut-9_F Frag389-436 61.7C | ATAGGTCTCCGAGACGtGCAACTAATTTTAGTCTAC |
| MOR_oligoP_DMS-1_F Frag2-49_55.5C | GACGTGTTACCCTCTCCTAG |
| MOR_oligoP_DMS-1_R Frag2-49_56.7C | TCGGACTCTCCCGTAGTAG |
| MOR_oligoP_DMS-2_F Frag50-100_58.0C | CAACGAGGTTTATTCGGAGG |
| MOR_oligoP_DMS-2_R Frag50-100_57.3C | TTACTCACGCGATGAATGGG |
| MOR_oligoP_DMS-3_F Frag101-150_57.3C | TAGAGCTATTCCCGGTTGAGG |
| MOR_oligoP_DMS-3_R Frag101-150_56.4C | GACTACACCTGTTATACCTCGG |
| MOR_oligoP_DMS-4_F Frag151-200_58.0C | TCTAGGTTTCGGCTTCATGGG |
| MOR_oligoP_DMS-4_R Frag151-200_55.9C | AGGGTGACTAGCGGTCAC |
| MOR_oligoP_DMS-5_F Frag201-250_56.9C | TTTTATGCTCTGTGTTGGCGG |
| MOR_oligoP_DMS-5_R Frag201-250_58.9C | AATGAGACTCGATACAGAGGGG |
| MOR_oligoP_DMS-6_F Frag251-300_58.6C | AATTGTGTTCGAGAAGTGCGG |
| MOR_oligoP_DMS-6_R Frag251-300_59.2C | CTCGTACCTCTATGATTGTTGG |
| MOR_oligoP_DMS-7_F Frag301-350_56.8C | ACCCAAAGAACTCGATTCCGG |
| MOR_oligoP_DMS-7_R Frag301-350_57.4C | TCTGTTCCTGAGTATTTGACGG |
| MOR_oligoP_DMS-8_F Frag351-400_56.9C | CTGAGACGATCCCTATCGAGG |
| MOR_oligoP_DMS-8_R Frag351-400_56.5C | GATACAAGGGAGGAATGATCGG |
| MOR_geneP_Mut-1_R Frag2-49 61.4C | ATAGGTCTCTCGATgctggcgtcatcatcg |
| MOR_geneP_Mut-1_F Frag2-49 60.5C | ATAGGTCTCacctcggtggtcgagatagc |
| MOR_geneP_Mut-2_R Frag50-100 56.6C | ATAGGTCTCttagggccacaagggtcg |

| | |
|---|---|
| MOR_geneP_Mut-2_F Frag50-100 59.7C | ATAGGTCTCaaaccgcaaccaatatttacatctttaatc |
| MOR_geneP_Mut-3_R Frag101-150 59.8C | ATAGGTCTCgtgtatctgacgataacgtacatgac |
| MOR_geneP_Mut-3_F Frag101-150 58.0C | ATAGGTCTCacatgttcacctccatttttaccttg |
| MOR_geneP_Mut-4_R Frag151-200 60.5C | ATAGGTCTCtcaattgagattacgattttgcacagaatg |
| MOR_geneP_Mut-4_F Frag151-200 57.8C | ATAGGTCTCtgccggtgatgttcatggc |
| MOR_geneP_Mut-5_R Frag201-250 61.0C | ATAGGTCTCgcgctagacagaatccaattgca |
| MOR_geneP_Mut-5_F Frag201-250 61.1C | ATAGGTCTCtttgttatggtcttatgatcctgcgc |
| MOR_geneP_Mut-6_R Frag251-300 58.8C | ATAGGTCTCatcagtactggcatgatgaaggc |
| MOR_geneP_Mut-6_F Frag251-300 56.9C | ATAGGTCTCttataattaaggccctggtgacc |
| MOR_geneP_Mut-7_R Frag301-350 56.7C | ATAGGTCTCtgaataggggtccagcaaaca |
| MOR_geneP_Mut-7_F Frag301-350 59.8C | ATAGGTCTCtctgcattcccacaagctctaac |
| MOR_geneP_Mut-8_R Frag351-400 61.9C | ATAGGTCTCaagcatcgcttgaaattctcgtcc |
| MOR_geneP_Mut-8_F Frag351-400 61.7C | ATAGGTCTCCGAGACGtGCAACTAATTTTAGTCTAC |
| VatD_oligoP_DMS-1_F Frag2-41_56.9C | TGGCATGCGCACTTACTG |
| VatD_oligoP_DMS-1_R Frag2-41_55.8C | CGTACCTTACGCATCCCAT |
| VatD_oligoP_DMS-2_F Frag42-84_56.3C | GTAAAACACAGGCCGCCAT |
| VatD_oligoP_DMS-2_R Frag42-84_56.4C | CTCCACGGTTTGTACAGTGTT |
| VatD_oligoP_DMS-3_F Frag85-126_56.7C | CACTACTCTTAGCCCGCTC |
| VatD_oligoP_DMS-3_R Frag85-126_58.4C | TGCGGCTGTTACGTTCAGT |
| VatD_oligoP_DMS-4_F Frag127-168_59.3C | ATCTCGTTTTCTCGCTCGGT |
| VatD_oligoP_DMS-4_R Frag127-168_57.0C | CCGTTATCTAGCTACGGCC |
| VatD_oligoP_DMS-5_F Frag169-209_58.2C | CACATCCTCGCTCGGTGTA |
| VatD_oligoP_DMS-5_R Frag169-209_56.7C | TAGTCCACCGCTACCCATG |
| VatD_geneP_Mut-1_R Frag2-41 57.9C | ATAGGTCTCGTAGGATCCGGTgtaagttatgtaa |
| VatD_geneP_Mut-1_F Frag2-41 61.1C | ATAGGTCTCGTGAGACATTTGACAAGCAAATCTTATACC |
| VatD_geneP_Mut-2_R Frag42-84 60.3C | ATAGGTCTCGAATCGTAATACGAATACTCACCT |
| VatD_geneP_Mut-2_F Frag42-84 59.3C | ATAGGTCTCGCTCAACCTACCCTTTTAATCTTTTTG |
| VatD_geneP_Mut-3_R Frag85-126 58.1C | ATAGGTCTCCGATGGTTTGCTCCATTCATAATGAT |
| VatD_geneP_Mut-3_F Frag85-126 57.9C | ATAGGTCTCTTATGCCCGGTGTAAAAATCGG |
| VatD_geneP_Mut-4_R Frag127-168 58.4C | ATAGGTCTCTCTTTTCCAATCCACACGTCATTAC |
| VatD_geneP_Mut-4_F Frag127-168 60.2C | ATAGGTCTCAGGACACTATCAATCAGTTACTTGAC |
| VatD_geneP_Mut-5_R Frag169-209 58.0C | ATAGGTCTCCGTTGTTTGATTTCATTTGCGGG |
| VatD_geneP_Mut-5_F Frag169-209 60.0C | ATAGGTCTCAGCTTGGTGactgcattctagttg |

Supplemental Table 3. **Primers used in this study.** Primer names and their sequences are listed that were used for amplifying DNA for sequencing and those that were used for generating DIMPLE libraries.

| Sample | Cells collected | Total Sequencing reads | Total variants | Filtered variants |
|---|---|---|---|---|
| Surface1_rep1 | 329590 | 20293422 | 1000418 | 561175 |
| Surface2_rep1 | 330554 | 20575732 | 953381 | 610207 |
| Surface3_rep1 | 630379 | 25184836 | 1166943 | 907847 |
| Surface4_rep1 | 494023 | 26135194 | 1163352 | 892634 |
| Surface1_rep2 | 600,000 | 18342202 | 827920 | 478076 |
| Surface2_rep2 | 450000 | 27550336 | 1364760 | 871004 |
| Surface3_rep2 | 1200000 | 23316806 | 1115989 | 870483 |
| Surface4_rep2 | 900000 | 24886520 | 1107320 | 858286 |
| Surface1_rep3 | 909976 | 33086780 | 1514613 | 1059664 |
| Surface2_rep3 | 947819 | 23558860 | 1117409 | 801760 |
| Surface3_rep3 | 978955 | 21732210 | 1017187 | 870483 |
| Surface4_rep3 | 1004080 | 22583636 | 1000418 | 829726 |
| Kir21 Baseline | >2000000 | 20996286 | 973733 | 750600 |
| TrpV1_1 | NA | 25377705 | 1357924 | 735263 |
| TrpV1_2 | NA | 14465421 | 904473 | 524229 |
| OPRM1 | NA | 1868121 | 134631 | 103943 |
| VatD | NA | 20361746 | 952787 | 745626 |

Supplemental Table 4. **Cell Sorting and Sequencing Summary Statistics.** Summary statistics for the number of cells collected per subpopulation for Kir2.1, total number of reads for each sample used in this study, number of reads that aligned to variants within the gene, and variants that we filtered for use in figures and fitness calculations because they were expected.

| Type | |
|---|---|
| **Deletions** | **58590** |
| 1 bp | 18224 |
| 2 bp | 1990 |
| 3 bp | 5478 |
| >3 bp | 32898 |
| **Insertions** | **8924** |
| 1 bp | 6908 |
| 2 bp | 1171 |
| 3 bp | 16 |
| >3 bp | 829 |
| **Synonymous** | **27963** |
| **Missense** | **30465** |
| **Other** | **52659** |
| **Total** | **178601** |
| **Designed variants** | **716056** |
| **Fraction Correct** | **0.8003693** |

Supplemental Table 5. **Baseline Kir2.1 library errors.** Error counts within the sequencing data. Because Illumina sequencing platforms have baseline error rates to a degree this is a combined error between sequencing and OLS DNA synthesis. Errors are broken up between deletions and insertions broken up across multiple length classes, point mutations that would result in synonymous and missense mutations, other which includes multiple mutations within a sequence which is a common within OLS subpool that many oligos with error will have multiple. Also included are the sum total and the number read counts of expected designed variants within the library. From this we can calculate that about 80% of our library are designed variants while 20% are not.
